# EXTRA-seq: a genome-integrated extended massively parallel reporter assay to quantify enhancer-promoter communication

**DOI:** 10.1101/2024.12.08.627402

**Authors:** Judith F. Kribelbauer-Swietek, Vincent Gardeux, Gerard Llimos-Aubach, Katerina Faltejskova, Julie Russeil, Nadia Grenningloh, Lucas Levassor, Clémence Steiner, Jiri Vondrasek, Bart Deplancke

**Affiliations:** Laboratory of Systems Biology and GeneKcs, InsKtute of Bioengineering, School of Life Sciences, École Polytechnique Fédérale de Lausanne (EPFL), Lausanne, Switzerland; Swiss InsKtute of BioinformaKcs, Lausanne, Switzerland; InsKtute of Organic Chemistry and Biochemistry, Czech Academy of Sciences, Prague, Czech Republic; Computer Science InsKtute, Faculty of MathemaKcs and Physics, Charles University, Prague, Czech Republic

## Abstract

Precise control of gene expression is essential for cellular function, but the mechanisms by which enhancers communicate with promoters to coordinate this process are not fully understood. While sequence-based deep learning models show promise in predicting enhancer-driven gene expression, experimental validation and human-interpretable mechanistic insights lag behind.

Here, we present **EXTRA-seq**, a novel **EXT**ended **R**eporter **A**ssay followed by **seq**uencing designed to quantify enhancer activity in endogenous contexts over kilobase-scale distances. We demonstrate that EXTRA-seq can be targeted to disease-relevant loci and captures expression changes at the resolution of individual transcription factor binding sites, enabling mechanistic discovery. Using engineered synthetic enhancer-promoter combinations, we reveal that the TATA-box acts as a dynamic range amplifier, modulating expression levels in function of enhancer strength. Importantly, we find that integrating state-of-the-art deep learning models with plasmid-based enhancer assays improves the prediction of gene expression as measured by EXTRA-seq. These findings open new avenues for predictive modeling and therapeutic applications.

Overall, our work provides a powerful experimental platform to interrogate the complex interplay between enhancers and promoters, bridging the gap between *in silico* predictions and human-interpretable biological mechanisms.

## Introduction

In eukaryotic genomes, enhancers and target promoters are often separated by large distances, ranging from hundreds of base pairs (bps) to hundreds of kilobases (kbs). How communication between a given pair of regulatory elements (REs) is established at a molecular level, and to what extent RE sequence determines enhancer-promoter (E-P) compatibility remains an open question^1^. Genome topology is thought to play a critical role, but contact frequencies alone are insufficient to predict enhancer-driven activity levels^2–7^.

Multiple experimental approaches have been used to uncover RE communication rules. For example, high resolution imaging has revealed the presence of enhancer ensembles (e.g. super enhancers^8^, condensates^9^, or more generally microenvironments^10^) whose assembly is mediated by coactivators, such as BRD4^11^, as well as disordered, or multivalent TF domains^12–14^. Complementary approaches include CRISPR-based enhancer perturbations^15,16^, or rely on epigenetic modification and gene expression profiling across genetically diverse populations^5,17–20^ or single cells^21^ to detect coordinated REs, defined as regions that covary in their epigenetic signatures. Genetic dissections of individual loci with coordinated REs have shown that enhancers can amplify each other’s activity, even when only lowly active on their own^22–24^. A recently developed three-enhancer reporter gene cell line has further shown that distance, and the order of strong and weak enhancers are important for cooperativity^25^. These orthogonal approaches have provided valuable insights into RE assembly and cooperativity. However, they have limitations in decoding the regulatory grammar that governs RE communication. Imaging studies are typically restricted to one regulatory architecture or TF at a time, while analyses of epigenetic coordination must infer mechanisms across diverse genomic contexts. This inference is particularly challenging given the complex network of possible interactions between regulatory proteins.

As an alternative to endogenous readouts, several episomal massively parallel reporter assays (MPRAs) have been developed to directly sample E-P, E-E, and even E-E-P interactions in high throughput^26–30^. In contrast to select genetic dissections, paired E-E activity was largely additive^30^ in episomal MPRAs. Independently, several studies have revealed that enhancers interacting with housekeeping *versus* developmental promoters deploy different regulatory programs, with multiplicative or supra-additive activity being more prevalent in promoters belonging to the latter group^27,30^. However, it remains unclear to what extent these plasmid-based assays faithfully reflect endogenous E-P communication. Developing methods that probe the molecular rules underlying native RE coordination is thus of high priority.

In parallel, the rapid advancement of deep learning-based tools has enabled the prediction of gene expression directly from sequence^31–34^. These models consider (several) hundreds of kbs, in principle allowing for inference of combinatorial regulatory mechanisms like E-P and E-E communication and cooperativity. Given this potential, several tools have been built on top of large sequence models to extract human-interpretable mechanisms from such ‘black box’ approaches^35,36^. Moreover, models like Enformer^31^ are trained on epigenetic and transcriptional data from hundreds of different cell lines, making them a suitable *in silico* validation option for generative AI tools. The latter aim to generate synthetic sequences with cell type-specific activity^37–42^, opening the door for targeted manipulations to the non-coding genome for therapeutic purposes. However, training data for such generative models are often derived from episomal assays^37,40,42,42^, or genome-wide readouts of local activity (e.g. DNA accessibility^39,41^ at the element itself), raising the question whether and to what extent synthesized sequences can truly substitute endogenous enhancers that require E-P communication across substantial distances.

Despite rapid computational advances, it has remained challenging to infer well-defined human-interpretable rules from deep learning models^43^ making it difficult to assess their ability to: capture the full complexity of enhancer-driven gene regulation, predict never-seen genetic contexts, and transfer to new cell types or species. Evidence from studies on personalized genomes suggests that enhancer-based gene regulation is still not sufficiently captured by existing models^44,45^.

To counteract the ‘black box’ approach in sequence-to-expression modeling, new experimental systems are required. These could not only enhance interpretability of existing models but may also improve their performance. Current experimental assays have a variety of shortcomings that limit mechanistic inference: i) they are not always genome-integrated, ii) they only test enhancers placed within short distances of a TSS, blurring the distinction between extended promoter versus actual enhancer activity^1,46^, or iii) they are confounded by different genomic contexts, i.e. each manipulation is embedded in a different enhancer context.

To overcome these challenges and provide deeper insights into the mechanistic underpinnings of commonly used experimental and computational tools, we developed a novel, genome-integrated and **ext**ended massively parallel **r**eporter **a**ssay followed by **seq**uencing. EXTRA-seq quantifies enhancer activity in a controlled genomic context by recording gene expression changes as a result of modifications in enhancers that are placed several kilobases away from the transcription start site (TSS). We demonstrate that EXTRA-seq can be targeted to endogenous loci of interest, that it enables probing combinatorial manipulations to both enhancers and promoters, and that it captures expression changes at the resolution of individual transcription factor binding sites (TFBS) required for mechanistic discovery.

By systematically comparing endogenous enhancer activity as measured by EXTRA-seq to the predictions of state-of-the-art deep learning models and traditional plasmid-based assays, we find that both data sources capture different aspects of enhancer function and that their integration improves gene expression predictions.

## Results

### EXTRA-seq workflow

To probe E-P communication across several kbs with the intent of more closely mimicking endogenous settings, EXTRA-seq relies on Cre recombinase-mediated cassette exchange (RMCE), as has previously been used to integrate short MPRA sequences (< few hundred basepairs^47–49^). To allow targeting a specific locus of interest (LOI), we used CRISPR-based cell engineering with two guide RNAs to insert a mono-allelic landing pad containing a CMV-GFP marker flanked by asymmetric lox sites (**Figure 1a**). To allow quantitative readouts of gene expression, the landing pad requires placement within the 5’ UTR of a haplosufficient gene, replacing the TSS, part of the UTR and a substantial part of the upstream regulatory region (i.e., a few kbs). This allows for the reassembly of the gene regulatory context under various perturbation settings, including modifications to both enhancer and promoter sequences (see **Methods** for details on the library assembly process). To track and monitor the impact of distant E-P modifications, each replacement sequence is multiplexed with random barcodes (BCs) that are embedded within the 5’UTR (**Figure 1a** and **Supplemental Figure 1a**). Libraries also contain a BC-upstream exogenous primer site for library amplification, as well as the two asymmetric lox sites on either end. ‘BC to modification’ mapping is achieved using long-read sequencing on the final plasmid pool (**Figure 1a,b**). Library-integrated cells are then selected based on loss of the GFP-cassette, with integration efficiencies ranging between 1-6% (**Supplemental Figure 1b)**.

**Figure 1.**
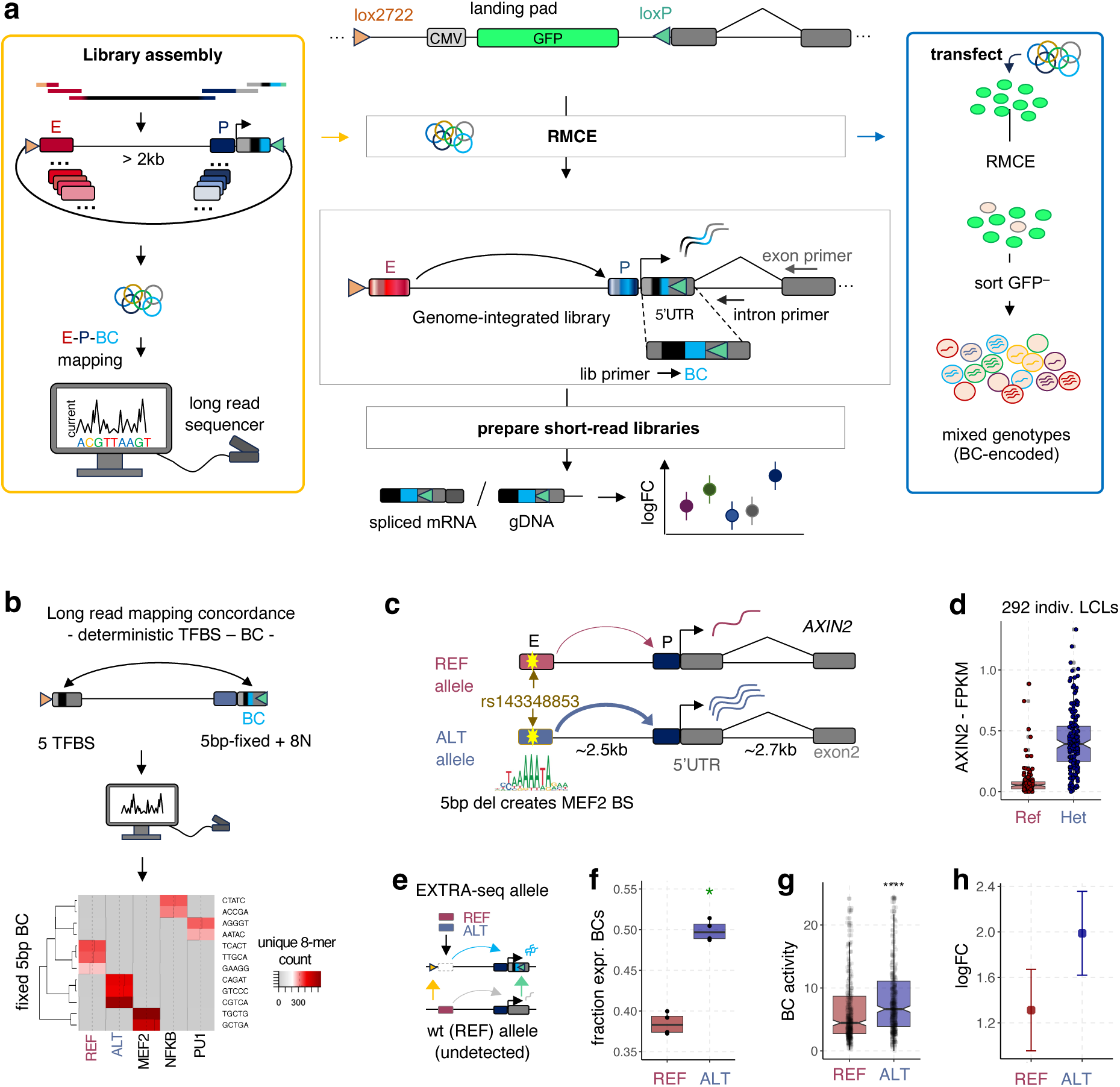
EXTRA-seq method and validation. **a** Overview of the EXTRA-seq approach. **Middle:** A lox-based landing pad is integrated at a focal location, replacing part of the 5’ UTR, the TSS, and the TSS-upstream regulatory region of interest (middle, top). The landing pad is replaced with a library encoding different enhancer (E) and promoter (P) modifications with unique barcodes (BCs) in the UTR. mRNA and gDNA are amplified using a fixed library (lib) and either exon-(mRNA) or intron-(gDNA) specific primers. The log fold change (logFC) of mRNA over gDNA ratios reflects construct-level expression. **Left**: Strategy to synthesize large, barcoded libraries with simultaneous modification to enhancers and promoters, and to map modifications to each BC. **Right**: Experimental strategy illustrating the use of BCs to capture pooled genotypes. **b** Validating mapping accuracy using a plasmid library containing pre-determined, enhancer-centric transcription factor binding site (TFBS)-to-BC associations. BCs contain a fixed TFBS-specific (5bps) and a random (8bps) part. Number of unique 8mers per fixed 5mer is shown across TFBSs. **c** Description of the rs143348853 variant locus at the *AXIN2* gene. A 5bp deletion creates a new MEF2 TFBS in the alternative (ALT) genotype enhancing *AXIN2* expression. **d** *AXIN2* expression levels (FPKM = fragments per kilobase of transcript per million mapped reads) across individually derived LCL cell lines from human donors split by their genotype for rs143348853. Homozygous reference (REF; no deletion) and heterozygous (REF/ALT) LCLs are shown to illustrate expected expression changes due to one copy of the ALT genotype. **e** Illustration of the *AXIN2* locus after placement of the EXTRA-seq cassette. **f** The fractions of ‘expressed’ BCs per replicate (non-zero counts in the mRNA library) are shown for either the REF or ALT EXTRA-seq allele (Mann Whitney test; p-value = 0.029). **g** Raw EXTRA-seq activities (averaged across replicates) per BC grouped by genotype (Mann Whitney test; p-value = 9.1*10^− 9^). Boxplots show the mean and quartiles with whiskers extending to 1.5 times the interquartile range. **h** Summarized log fold change (logFC) in expression using the mpralm function^50,51^. Whiskers indicate the scaled standard deviations associated with each fit.

To obtain quantitative readouts of expression without the need to establish individual cell lines, we simultaneously extract genomic (g) DNA and mRNA for a minimum of three replicates from each GFP-negative, and library-integrated cell pool. The barcode-containing 5’UTR region is then amplified using either an intron-(for gDNA amplification), or an exon-(for mRNA amplification) specific primer (**Figure 1a**). after which enhancer-driven gene expression for each modification is computed by taking the ratio of mRNA over gDNA BC counts.

### EXTRA-seq captures expression changes driven by enhancer-centric non-coding variation

To demonstrate the feasibility of using EXTRA-seq in a biomedicine-relevant context, we focused on the *AXIN2* gene, as its expression is regulated by an approximately ∼2.5kb distant enhancer that contains a naturally occurring, leukemia-protective 5 bp deletion (rs143348853). The deletion results in the recruitment of the TF MEF2, which in turn enhances *AXIN2* expression^2^ in individuals with the alternative (ALT) genotype (**Figure 1c,d**). To integrate the EXTRA-seq landing pad within one allele of the *AXIN2* locus, we designed guide RNAs targeting the 3’ end of the indel-containing enhancer and the first exon of *AXIN2* (5’UTR; ∼100bp downstream of the TSS, **Supplemental Figure 1a**) in the chronic lymphocytic leukemia (CLL) cell line MEC1, whose genotype is homozygous reference for rs143348853.

To explore whether EXTRA-seq is able to restore endogenous gene regulatory architectures, we generated libraries that, when integrated, reestablish the rs143348853 reference (REF) or ALT genotypes (one copy, **Figure 1e**). Both genotypes were multiplexed with hundreds of random 10bp-long barcodes before integration into the modified *AXIN2* locus. As EXTRA-seq relies on pooled genotypes, we verified that independent, technical replicates show significant concordance at the level of both gDNA-and mRNA-based barcode counts, ensuring that each pool captures the full range of integrated barcodes (**Supplemental Figure 1c,d**). The proportion of BCs with zero counts was higher in the mRNA samples, suggesting that, while most BCs are reliably detected at the gDNA-level, not all barcodes lead to detectable expression levels. To test if the ALT genotype is indeed more active, we computed the number of ‘expressed’ (non-zero) BCs per replicate as a simple heuristic for expression differences between the two genotypes (**Figure 1f**). On average, the fraction of expressed BCs was 10% higher for the ALT genotype. Comparing the distributions of average, per-barcode activity levels (i.e. the ratio between mRNA and gDNA counts) between REF and ALT directly, we similarly found the ALT genotype to drive significantly higher expression levels (p-value = 9.1*10^−9^, Mann-Whitney test, **Figure 1g**).

Finally, we used the mpralm package^50,51^, specifically designed for MPRA-type assays to summarize the genotype-specific expression levels, revealing on average a roughly 1.6-fold expression difference between ALT and REF (**Figure 1h**). Together, these findings indicate that EXTRA-seq recapitulates the effect of rs143348853 in CLLs in heterozygous condiVons, not only validaVng our approach but also demonstraVng its ability to measure variant effect sizes.

### EXTRA-seq reveals large variability of strong enhancers in out-of-context settings

To assess EXTRA-seq’s ability to capture variability in E-P communication more generally, we designed a library containing REF, ALT, as well as different *AXIN2* enhancer replacements, including i) a complete enhancer deletion, ii) ten highly active B cell enhancers and five non-enhancers from genomic loci other than *AXIN2* (200 bp; for the selection strategy, see ^41^), and iii) three synthetic enhancers that contain three or four top binding sites of TFs with specific contributions to B cell enhancer function (i.e. MEF2, PU.1, FOX, NFkB and BACH)^52^. First, we verified that the inferred genotype-specific expression levels were highly reproducible across replicates and that there was no relationship between activity and integration frequency (**Supplemental Figure 2a,b**) after correcting for the mean-variance relationship using the mpralm package (**Methods**). Comparing enhancer-specific expression levels, we found that, as expected, neither the five non-enhancer sequences, nor the enhancer deletion was capable of activating *AXIN2* expression above the baseline level set by the REF enhancer (**Figure 2a**). In contrast, two of the three tested synthetic enhancers (each containing four distinct TFBSs) were among the genotypes driving the highest levels of expression, similar to the strongest endogenous enhancers (**Figure 2a**). Moreover, we observed a large degree of expression variability among B-cell enhancers, even though all had been selected based on high local activity in their native contexts. To investigate what drives this variability, we compared EXTRA-seq-measured expression to several native, molecular enhancer signatures, including DNA accessibility, histone modification levels, and coregulator recruitment (**Figure 2b**). Focusing on the ten active enhancers, we found that only H3K27ac and acetyl-lysine binding coactivator BRD4 enrichment levels, but not H3K4me1 or acetyltransferase binding (p300), are predictive of expression (Pearson’s rho = 0.71 (H3K27ac) and 0.76 (BRD4), p.val = 0.02 and 0.01, respectively). Interestingly, BRD4 levels, a coactivator we had previously linked to cooperative and communicating enhancer environments^52^, and which is also associated with super enhancer formation, correlated slightly better with expression than enhancer acetylation levels alone.

**Figure 2.**
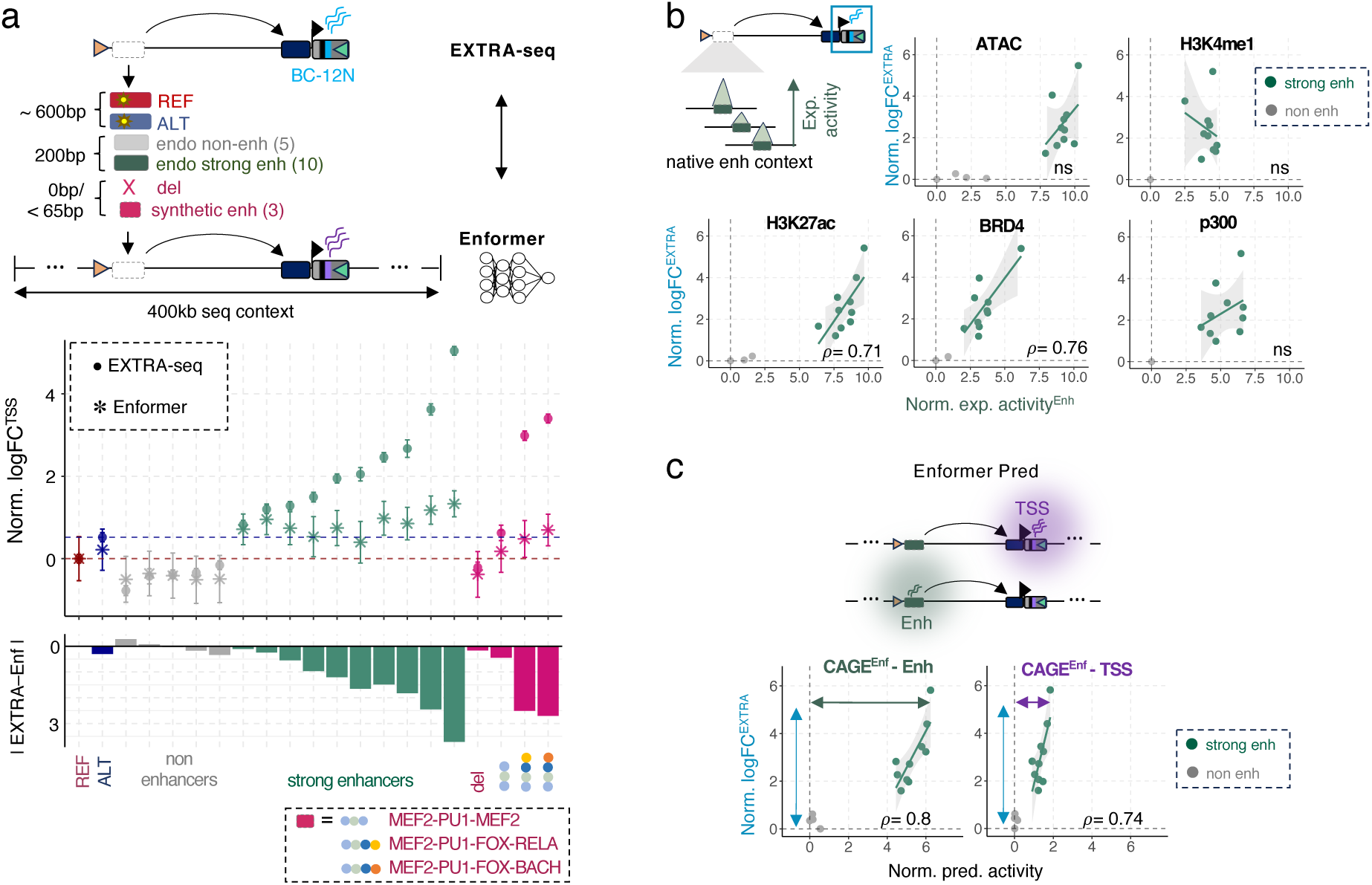
Out-of-context enhancers drive highly variable AXIN2 expression. **a** Overview of experimental strategy and EXTRA-seq library design (top), and results (bottom). Beyond the REF and ALT enhancer, several synthetic and endogenous (non-) enhancer fragments replace the endogenous *AXIN2* enhancer. Scatterplot shows logFC of EXTRA-seq derived (points) or Enformer-predicted (stars) expression levels normalized by the respective levels for the REF genotype. Error bars represent standard deviations associated with either experimental activities or computational predictions. Barplots show the absolute difference in normalized logFC between the measured activity and the predicted activity. Colors indicate different enhancer classes, with colored points representing individual TFBSs for synthetic enhancers. **b** EXTRA-seq logFC versus local enhancer activities (normalized by the fragment with lowest activity, respectively) for five different enhancer signatures as measured in the enhancer’s native context. Colors indicate strong (green) versus non (gray) enhancers. Lines and shaded areas represent linear fits for strong enhancers. Correlation levels are indicated for enhancer features that exhibit significant correlations. Exact p-values = 0.02 for H3K27ac, and 0.01 for BRD4. **c** Same as in **b** but using Enformer-predicted activity levels (CAGE^Enf^) at either the enhancer locally or the TSS/UTR for each of the 15 endogenous enhancer fragments. Exact p-values = 5.3*10^−3^ for CAGE-Enh, and 0.02 for CAGE-UTR.

Finally, given the immense interest in training computational models to predict gene expression, we compared EXTRA-seq expression levels to *in silico* predictions made by the widely used transformer-based deep neural network Enformer^31^. Of note, we spotted a potentially problematic behavior of Enformer-based predictions, with the predicted CAGE signal at the *AXIN2* TSS varying substantially as a function of subtle changes in the positional encoding of the replacement cassette within the input sequence (shifts of a few bps; **Supplemental Figure 3a**). We found that this variability translates into substantial effect size prediction differences for the REF/ALT genotypes (varying from no effect at all to effect sizes 3-times the average effect size; **Supplemental Figure 3a**), both at the *AXIN2* enhancer, as well as the promoter (**Supplemental Figure 3b**). To remedy this observed variability at the *AXIN2* locus, we averaged effect size predictions across several positional encodings (**Methods**). Using this strategy, we found that Enformer-based predictions of CAGE signal at the TSS tend to underpredict the observed range in EXTRA-seq expression across enhancers, with the discrepancy between measured and predicted activity increasing with enhancer strength (**Figure 2a**). Moreover, using BRD4 enrichment from the native enhancer context correlated better with the out-of-context EXTRA-seq expression levels than TSS-based predictions of CAGE (**Figure 2b,c**). Surprisingly, however, Enformer-based predictions of enhancer-centric CAGE levels not only showed stronger correlations with final expression levels, but also captured the full dynamic range (**Figure 2c**). Enhancer-centric predictions even outperformed native enhancer marks, supporting the notion that deep learning models effectively capture local activity, but face greater challenges in modeling RE communication^44,45^.

### EXTRA-seq enables the sensitive assessment of enhancer activity at nucleotide resolution

Our results so far demonstrated that EXTRA-seq can capture differences between sequences (endogenous and exogenous alike) encoding functional enhancers versus those that do not, and that, in principle, it can identify epigenetic signatures indicative of productive E-P communication. In a next step, we explored EXTRA-seq’s sensitivity to capture effect sizes of non-coding genetic variation at the level of single TFBSs. To this end, we tested a library of 19 individual TFBSs replacing the sequence spanning the critical MEF2 TFBS at the rs143348853 variant and tested their ability to activate *AXIN2* expression (**Figure 3A)**. We again included key B-cell TF motifs (IRF, BATF, NFkB, etc.), CTCF, as well as motif variations (spacing, flanking sequence, and copy number) for the TF PU1, a pioneer TF that is active in B cells^53^ and belongs to the group of ETS factors. In addition to EXTRA-seq, we also tested the REF/ALT enhancers, as well as all 19-TFBS enhancer variations, using two commonly used methods to measure or predict variant effect sizes: i) STARR-seq^54^ (representing plasmid-based MPRAs, **Methods**), and ii) Enformer (representing *in silico*-based approaches). Comparing activity levels across the 21 total enhancers, we found that several TFs other than MEF2 are capable of activating *AXIN2* expression, with two copies of the PU1 site achieving highest levels (**Figure 3a**). To assess the reproducibility of the derived activities for individual TFBSs, we compared EXTRA-seq activities obtained from eight TFBSs that were included in three independent library transfections (**Supplemental Figure 4a** and **Methods**). Albeit slightly more variable than the inter-replicate activities derived from the out-of-context enhancer libraries (**Supplemental Figure 2a**), we still obtained a high overall correlation (Pearson’s rho between 0.82 and 0.93, **Supplemental Figure 4b**), indicating EXTRA-seq’s ability to detect even subtle differences in expression.

**Figure 3.**
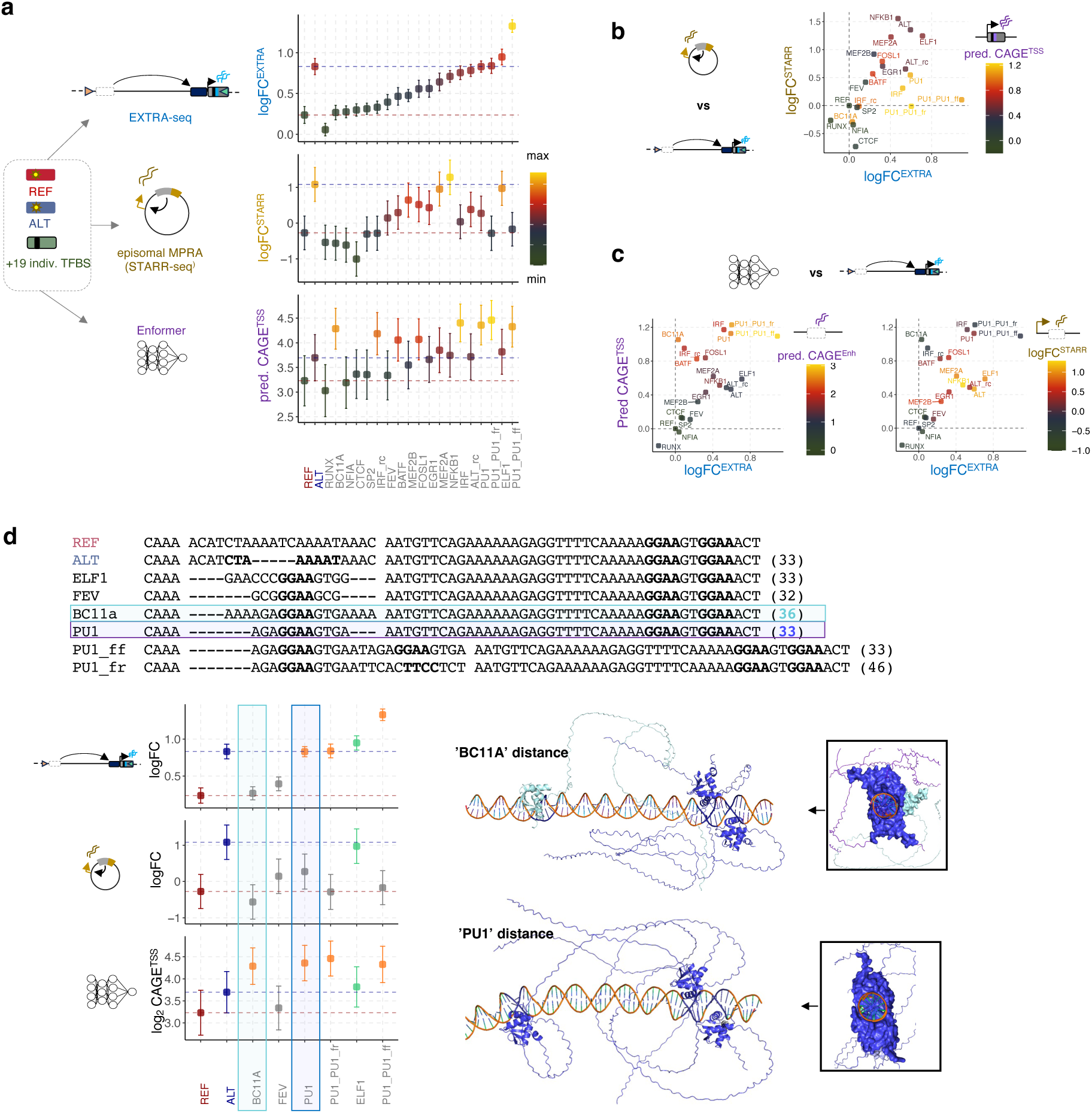
EXTRA-seq discovers expression differences of individual TFBS and BS configurations. **a** Experimental design and results. Left: In addition to the REF/ALT rs143348853 variant, several additional single- and dimeric TFBS genotypes are created and their variant-effect size measured using EXTRA-seq, STARR-seq or Enformer. Scatterplots show the measured or predicted expression levels (logFC or log predictions) associated with each genotype for each tool. Constructs are ordered by their effect size in the EXTRA-seq library, with REF and ALT genotypes indicated by red and blue dotted, horizontal lines. Colors are scaled to the minimum and maximum expression levels for each tool. Whiskers represent standard deviations of average activities associated with each genotype. **b** Tool comparison between STARR-seq and EXTRA-seq. Shown are logFC expression levels across tested genotypes. Colors indicate Enformer-predicted expression levels at the TSS. **c** Same as in b but comparing Enformer-predicted (CAGE at the TSS) to EXTRA-seq expression. Color indicates the predicted activity at the variant-enhancer (CAGE^Enh^, left), or the measured activity using STARR-seq (right). **d** Comparison of tools for ETS TFBS variations. ETS-core sequences (GGAA) are highlighted in bold. Scatterplots show disinct differences across tools (orange = Enformer predicted, and green = STARR-seq predicted). Purple and blue colored boxes highlight the two ETS-sites with differential activity between Enformer and EXTRA-seq, despite their BSs being identical. Bottom right shows Alphafold3-predicted structures for those two sites. Boxes show views along the DNA axis.

When comparing to both episomal and *in silico* approaches, we found differences both in terms of ranking, as well as the magnitude of effect sizes (**Figure 3a**). Importantly, discrepancies to EXTRA-seq-derived variant effect sizes were approach-specific: while STARR-seq captured the expression increase associated with the original variant (REF/ALT) and activator TFs such as NFkB (**Figure 3b**), it discounted TFs linked to increases in DNA accessibility and predicted enhancer activity such as PU1 (**Figure 3b & Supplemental Figure 4c**). This aligns with prior observations that episomal assays tend to discount the activity of TFs linked to DNA access^52^. Enformer, by contrast, successfully captured these ‘accessibility-linked’ activities but tended to overestimate their effect sizes (**Figure 3b,c**).

Since the endogenous *AXIN2* enhancer already contains a tandem PU1 site within <35bp of the variant, we next focused on the 6 enhancers that contained one or more ETS factor TFBSs (all containing the ‘GGAA’ ETS core, **Figure 3d**). We found that four out of six drive expression in the EXTRA-seq setup (**Figure 3d**). STARR-seq captured the activity of one (‘ELF’; TFBS, unique flank) of those four TFBSs, indicating that the canonical ‘ELF’ site may preferentially recruit an “activator” ETS-TF and not the ‘pioneer’ PU1. All tools agreed that the ‘FEV’ site, which similar to the ‘ELF’ one contains a unique GGAA-flank, does not activate the enhancer. Enformer captured all four TFBSs active in EXTRA-seq. However, it also predicted activity for a fifth one, and did not differentiate among the ‘tandem PU1’ sites with different protein orientations. When focusing on the fifth motif predicted by Enformer (corresponding to a ‘BC11A’ site), we noticed that it had the exact same core and flanking sequence as the canonical ‘PU1’ site (AGA**GGAA**GTG). The only difference is a short 3-bp DNA spacer (**Figure 3d**, colored boxes), which increases the distance of the ‘GGAA’ core to the tandem PU1 sites already present in the native enhancer (**Figure 3d**).

To investigate whether configurational differences and protein positioning may explain the lack of activity for the ‘BC11A’ enhancer in EXTRA-seq, we predicted structures for both configurations using AlphaFold3^55^. We found that the predicted structure of the ‘PU1’ motif (shorter spacer) places the third PU1 protein on the same DNA face as one of the ‘dimer’ PU1s, resulting in substantial DNA bending. In contrast, the ‘BC11A’ motif (longer spacer) was predicted to result in a perpendicular arrangement off all three PU1 proteins, with the deformative forces exerted by each protein upon DNA engagement seemingly acting independent of each other (**Figure 3d**). Similar to previous findings, these results suggest that differences in protein alignment^49^ and TFBS positioning^56–58^, as well as the resulting differences in DNA bending may play a role in a TF’s ability to activate an enhancer, thus indicating that EXTRA-seq is capable of capturing changes in activity down to the level of distinct TF configurations. However, further experimental evidence will be required to ascertain whether in the context of the *AXIN2* enhancer, different protein alignments may indeed influence PU1’s ability to pioneer nucleosome-bound DNA, thus leading to preferential activation.

### EXTRA-seq reveals intricate regulatory promoter architectures not captured by ML models

To illustrate the versatility of EXTRA-seq, we next applied it to measure gene expression changes caused by simultaneous enhancer-promoter modifications. For this, we designed libraries containing nine different promoter modifications in combination with four different enhancer variations, including the REF, ALT, as well as the aforementioned PU1 and NFkB enhancers. The native *AXIN2* promoter includes an INR/DPE motif combination, and contains both SP2 and NFY motifs upstream of the TSS, which are thought to induce transcription in a bidirectional manner^33^. To cover a large variety of promoter motif functions, we made mutations abrogating either the INR, or the DPE motif, explored a total of six SP2 motifs, inversed the NFY-motif (bidirectionality test), and inserted a TATA-box (or a TATA-BREu motif combination), which are missing in the wildtype promoter. In addition, we deleted a ∼ 250bp G/C-rich stretch (including some of the SP2 motifs and extending further upstream) and paired all variations including the wildtype promoter with the four different enhancer types (**Figure 4a**).

**Figure 4.**
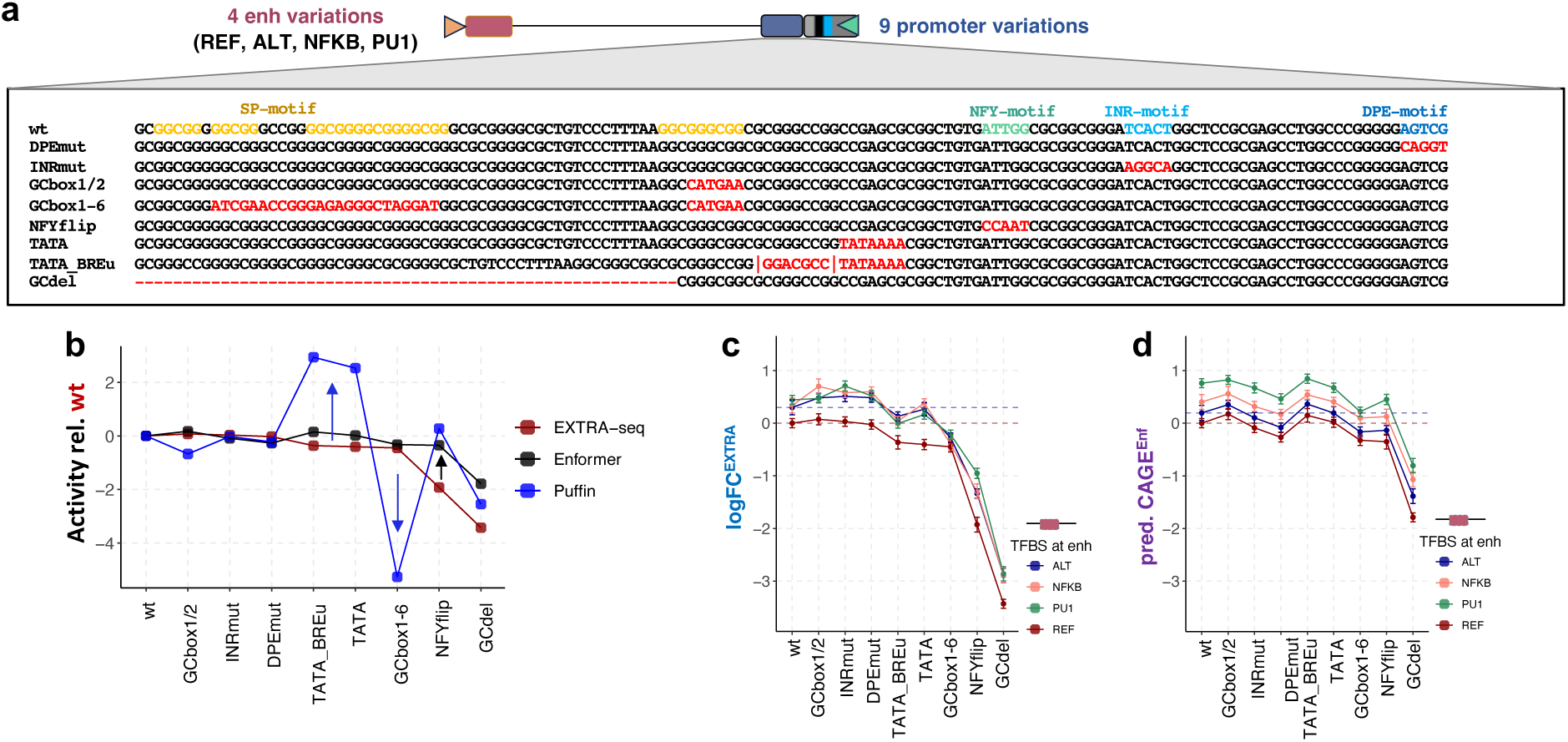
Promoter core motif function is highly context-specific. **a** Library design and promoter sequences for the nine different *AXIN2* promoter modifications. Core motifs indicated with different colors in the wildtype sequence, with red font highlighting individual modifications. Red dashed line indicates the start of the GC-deletion. **b** Tool comparison of promoter activities (REF enhancer setting) normalized relative to the activity of the wildtype promoter sequence. Different tools indicated by different colors. Arrows indicate substantial deviations. **c** EXTRA-seq expression logFC as a function of both promoter variations (x-axis) and different enhancer genotypes (colored lines) normalized by the wildtype promoter x REF enhancer construct. Red and blue dotted lines indicate REF and ALT genotypes. **d** Same as in c but for Enformer expression predictions.

First, we only used the ‘inactive’ REF enhancer to assess promoter motif effects in isolation. To our surprise, EXTRA-seq-measured promoter activities aligned poorly with the expectations set by the recently developed deep neural network model Puffin, which we chose for its interpretability (**Supplemental Figure 5**). For instance, out of the four promoter variations predicted by Puffin to have the largest effects, only one had a substantial effect in the same direction in EXTRA-seq. In contrast to the predictions, deletions of a couple, or even a handful of SP2 motifs only had mild expression effects, while the G/C-rich stretch deletion (GC-del) had a strong detrimental impact that goes beyond the deleterious effect of removing four SP2 motifs which drives Puffin’s prediction (**Figure 4b**). In addition, we did not observe the expected boost in expression when introducing a TATA-box. Strikingly, the inversion of the NFY motif, which is thought to be bidirectional, had the second strongest deleterious impact on baseline expression. Given the striking differences with the Puffin model, we also included Enformer-based predictions, which we found to correspond better in that they did not predict large effects where none were observed by EXTRA-seq. Nonetheless, Enformer also failed to predict the deleterious effect of the NFY-motif inversion. These findings highlight that intricate regulatory architectures at individual gene promoters are poorly captured by models, whose performance is benchmarked against ‘average’ effects, thus demonstrating the need for genome-integrated tools to hone in on individual contexts.

Finally, we compared all E-P combinations as measured by EXTRA-seq and predicted by Enformer (**Figure 4c,d**). While all three non-REF enhancers (ALT = MEF2, PU1, NFKB) drove expression to similar degrees (thus confirming our previous EXTRA-seq findings, i.e. compare to **Figure 3a**), Enformer stratified motifs and favored PU1. Importantly, the effect size difference between enhancers was uniform across different promoter variations in Enformer, while our experimental EXTRA-seq read-outs were more nuanced. Notably, we observed a reduction in expression output in the promoter with a TATA-box-only insertion compared to wildtype for the REF enhancer, while the reduction was not observed for the other three, more active enhancers (**Figure 4c**).

### EXTRA-seq reveals nonlinear behavior between promoter architecture and synthetic enhancer strength

Our assessment of a limited number of E-P combinations provided interesting insights into the regulatory impact of promoter motifs in function of enhancer type, motivating the design of a larger library. The latter included REF/ALT as controls, 47 synthetic enhancer sequences of 1-5 TFBSs as well as a complete enhancer deletion, all of which were then paired with either the wildtype, TATA-box or GC-del promoters (**Figure 5a**). We observed a wide range of expression levels (∼ 200-fold change) across tested combinations, with a 16-fold range for the wildtype promoter alone (**Figure 5b).** To verify the reproducibility of the obtained expression levels, we sampled a smaller number of wildtype promoter constructs (∼30) and created a separately synthesized, cloned, and transfected library. Expression levels between the full and subsampled libraries strongly correlated (**Supplemental Figure 6a**; Pearson’s rho of 0.98, p-value < 0.0001) suggesting that EXTRA-seq produces consistent expression measures even for entirely synthetic enhancer constructs. Comparing between either ‘wildtype (INR-only) and TATA-box’, or ‘wildtype and GC-del’ promoters, we found a general correlation across enhancers (**Supplemental Figure 6b**), consistent with previous findings indicating that different regulatory environments may simply scale activities of promoters with intrinsic strengths^48^. Next, we probed the relationship between expression output and the number of TFBSs within an enhancer, and found that EXTRA-seq activity responds nonlinearly to the number of TFBSs for both wildtype and TATA-box promoters (**Figure 5c**), likely reflecting thresholding behavior induced by different TF occupancy levels, as described previously^59^. Importantly, this effect was not simply due to differences in homotypic versus heterotypic TFBS enhancers (**Supplemental Figure 6c**). Indeed, while this nonlinear trend was observed overall, individual TFBS combinations still varied drastically (**Figure 5c, and Supplemental Figure 6c**; see the ‘three TFBS’ enhancers).

**Figure 5.**
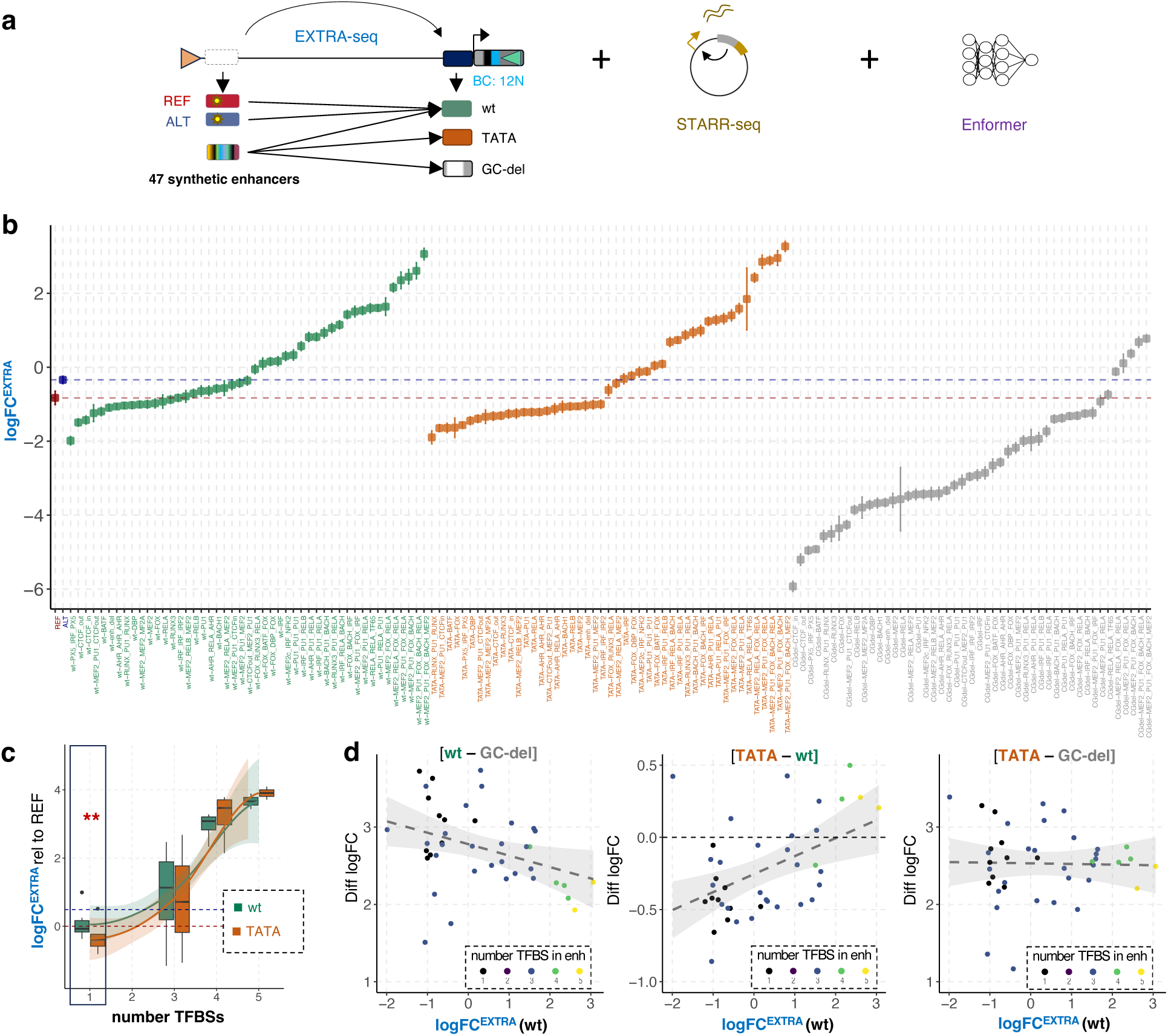
Core promoter elements interpret enhancer input in a strenth-dependent manner. **a** Library and method overview. **b** EXTRA-seq expression logFC across all tested genotypes. Colors indicate different promoter variations (green = wildtype (wt), orange = TATA-box, grey = G/C-rich stretch deletion or GC-del). **c** EXTRA-seq expression logFC of synthetic enhancer constructs normalized by the REF genotype as a function of TFBS number. Colors indicate different promoter variations (green = wildtype, orange = TATA-box). Black box shows significant differences between promoter variations (Mann-Whitney test; p-value = 2.8*10^−3^). **d** Pairwise differences in EXTRA-seq expression logFC across promoter variations in function of enhancer strength (logFC in wildtype promoter setting). Colors indicate the number of unique TFBSs across synthetic enhancers. Dashed lines and shades represent linear fits.

### EXTRA-seq reveals the TATA-box as a dynamic range amplifier

While activities between TATA-box and wildtype promoters were highly correlated across enhancers, we nonetheless noticed subtle differences in the absolute range of expression (**Figure 5b**). When splitting enhancers based on the number of TFBSs, we found that TATA-box promoters significantly lowered expression output for 1 TFBS constructs (p-value = 2.8*10^−3^, Mann-Whitney test, **Figure 5c**). This finding is not consistent with Enformer-derived predictions (**Supplemental Figure 6d**) and more generally, with the notion that the TATA box exclusively serves as a transcriptional amplifier. This effect disappeared for ‘three TFBS’ enhancers and seemingly inversed for ‘four’ and ‘five TFBS’ enhancers. The lowered expression for ‘weak’ (one TFBS) enhancers is in line with the observation already made in our library probing 9 different promoter sequences (see **Figure 4c**), namely that the TATA-box insertion lowered expression of the less active (i.e. weak) REF enhancer, while the three more active enhancers remained unchanged.

To investigate this phenomenon in more detail, we stratified the pairwise difference in logFC across all three promoter modifications (wildtype, TATA-box, and GCdel) by the wildtype enhancer strengths as measured directly by EXTRA-seq (**Figure 5d**). This analysis confirmed that the TATA-box switches from lowering expression for weak enhancers to amplifying expression for strong enhancers, thus enlarging the dynamic range (Pearson’s rho = 0.51, p-value =5.8*10^−4^, Figure 5d). Intriguingly, our analyses also revealed a dependency for the GC-del promoter on enhancer strength: while the GC-stretch boosted expression for all enhancers, the boost was more prominent for weak enhancers (Pearson’s rho = –0.36, p-value = 0.02). When comparing between the TATA-box and the GC-del promoter, we observed that the boosting due to the GC-stretch cancels out the ‘noise suppression’ effect on weak enhancers exerted by the TATA-box (**Figure 5d**).

### Combining local predictions and experimental measures of enhancer activity improves gene expression predictions

Given that we tested ∼ 50 different synthetic enhancers, we next asked whether EXTRA-seq-derived gene expression measures can be leveraged to improve gene predictions based on conventional tools by serving as an orthogonal, and, importantly, genome-integrated benchmark. To do so, we generated STARR-seq activities, as well as Enformer-derived predictions (wildtype promoter, including CAGE, enhancer accessibility (DNase), and enhancer activity (H3K27ac) enrichment levels) for all enhancers. Next, we systematically compared the correlation between EXTRA-seq enhancer activities to either STARR-seq activities, or predictions made by Enformer for both the TSS (CAGE) and the enhancer (CAGE, DNase, H3K27ac) (**Figure 6a**). Similar to our results for the ‘out-of-context’ enhancers, we found that Enformer predictions at the TSS did slightly worse than using predictions for the enhancer itself, in particular when considering the predicted expression range (**Figure 6a**).

**Figure 6.**
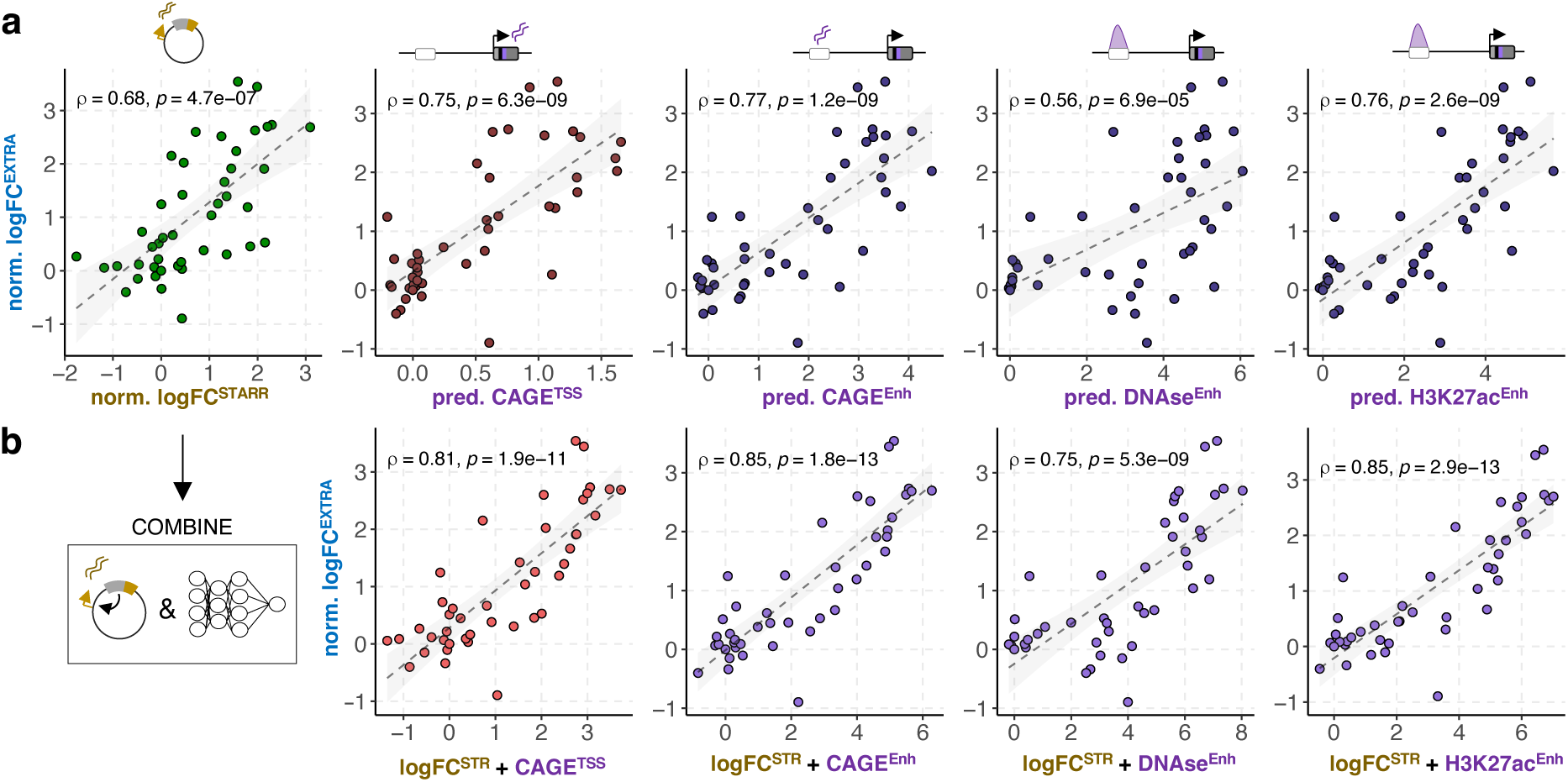
EXTRA-seq benchmarks new gene prediction strategies. **a** EXTRA-seq expression logFC relative to enhancer deletion genotype for 47 synthetic enhancers paired with the wildtype promoter in function of either STARR-seq logFC, or Enformer-predicted expression (CAGE^TSS^), enhancer expression (CAGE^Enh^), enhancer accessibility (DNAse^Enh^), and enhancer activity (H3K27ac^Enh^). Dotted lines and shades represent linear fits. Pearson’s rho and exact p-values are given in each plot. Colors indicate different tools (green = STARR-seq, red = Enformer) or different locations of activity asessment (Enformer prediction at TSS = red, at enhancer = purple). **b** Same as in **a** but combining STARR-seq end Enformer-predicted activities (sum of log-scaled predictions and logFC activites).

STARR-seq activities, albeit showing a significant correlation performed slightly worse than Enformer (Pearson’s rho = 0.68 versus 0.77, **Figure 6a**). Nonetheless, since our analysis of individual TFBSs had shown that STARR-seq and Enformer appear to capture distinct aspects of TF and thus enhancer function (i.e. STARR-seq captures TFs associated with intrinsic activity, while Enformer highlights those linked to DNA accessibility and cell type identity), we explored whether combining both tools would provide a better estimate of an enhancer’s endogenous function than each tool on its own. We found that integrating activities derived from STARR-seq with the different Enformer predictions resulted in improved correlations with EXTRA-seq expression across the board (**Figure 6b**). We observed the largest increase when combining STARR-seq activity with predictions for enhancer accessibility (increase in Pearson’s rho from 0.56 to 0.75), while the best correlation was obtained for the integration of STARR-seq and predicted enhancer activity (**Figure 6b**; enhancer CAGE: Pearson’s rho = 0.85, p-value = 1.8*10^−13^; enhancer H3K27ac: Pearson’s rho = 0.85, p-value = 2.9*10^−13^). These results reinforce the idea that both tools capture different aspects of enhancer function, and that their integration could serve as a valuable strategy for improving gene expression predictions.

## Discussion

We present EXTRA-seq, a genome-integrated assay that enables the analysis of enhancer-promoter communication over kilobase-scale distances, providing a powerful tool to dissect the molecular mechanisms underlying endogenous enhancer-driven gene regulation. Among EXTRA-seq’s key features are i) its integration into a controlled genomic context, which is useful for determining which chromatin signatures provide the best prediction for enhancer-promoter communication and downstream expression; ii) its ability to be targeted to specific loci, allowing for a detailed investigation of non-coding variants or disease-linked mutations in their native environment; iii) its use of barcodes, enabling a highly quantitative readout, effectively replacing the need to establish individual cell lines for each tested genotype; and iv) its high sensitivity, detecting quantitative differences in enhancer-driven gene expression even at very low expression levels (e.g. as observed for the GCdel promoter). This sensitivity allows for the resolution of expression differences at the level of individual TFBSs and even TFBS configurations, paving the way for screening insulator and repressor sequences in the future.

EXTRA-seq also allowed us to uncover a new regulatory feature intrinsic to the TATA-box motif: it acts as a dynamic range amplifier, suppressing noise from weak enhancers while amplifying strong enhancer activity. While its ‘amplification’ role has been recognized^60^, this ‘noise suppression’ is to our knowledge a new feature. We speculate that the latter function has been missed due to the challenge of observing decreased expression of already lowly expressed genes in RNA-seq data. In developmental contexts, this mechanism may be advantageous to restrict and tune expression: when important TFs are missing and enhancer strength is weak, the TATA-box prevents spurious activation. When enhancers gain strength with newly occupied TFBSs, the TATA-box switches to its known role as an ‘amplifier’. We observed similar, yet opposing dependencies on enhancer strength for the GC-rich stretch, suggesting that promoters use several licensing mechanisms to interpret enhancer inputs. Our findings also align with a previous episomal enhancer-promoter interaction study that showed that weaker promoters are more susceptible to ‘enhancer-enhancer boosting’^30^. Here, we propose that this phenomenon may generally hold true for enhancer-promoter interactions and that its magnitude and effect size may depend on specific promoter motifs. Thus, our findings suggest a bidirectional regulatory relationship between enhancers and promoters. Future studies could explore the generalizability of the observed effects, resolving how different promoter motifs interact and which molecular mechanisms underlie the differential interpretation of enhancer strength.

By systematically comparing EXTRA-seq with widely used experimental and computational variant effect size-estimation tools, we revealed method-specific biases. Computational tools struggled to capture subtle, but important changes in motif configurations (e.g. PU1, or NFY motifs), while STARR-seq was unable to capture ‘pure’ pioneering activity. More generally, while deep learning-based approaches appear well-suited for estimating the effects for thousands of variants, they appear less reliable when applied to a specific locus of interest, consistent with previous findings^44,45^. EXTRA-seq addresses these limitations by providing a near-native, yet controlled enhancer-promoter interaction environment, which serves as an orthogonal benchmark for ‘endogenous’ enhancer activity. Using this strategy, we found that computational tools and plasmid-based MPRAs complement each other in characterizing enhancer function. We therefore believe that EXTRA-seq offers opportunities to improve predictive algorithms (e.g. by systematically including episomal readouts), and explore novel applications in synthetic biology and therapeutics. Specifically, EXTRA-seq could enable the design of enhancer-promoter interaction systems that not only drive cell type-specific gene expression, but also achieve precise, pre-determined expression levels.

While EXTRA-seq represents an advancement over existing methods, it also has several limitations. First, since libraries are genome-integrated, the number of testable fragments is currently limited. This ranges from a few dozen (when probing individual TFBSs with relatively small effects, thus requiring >100 individual BCs) to a few hundred (when testing different enhancers with larger expected expression ranges, requiring fewer barcodes) fragments. A key constraint is the recombination efficiency in a given cell type. However, recent improvements in efficient and minimally scarring payload integration^61^ are expected to significantly enhance throughput. Second, the enhancer-promoter distance is limited by the logarithmic decrease in cloning efficiency for longer fragments, constraining the study of long-range (e.g. > 20kb) enhancer-promoter communication with this method. Finally, while EXTRA-seq provides relative comparisons between constructs, our results are specific to the *AXIN2* locus. Expanding our approach to more contexts and cell types and deriving a universal EXTRA-seq cassette will be valuable next steps to enhance generalizability.

## Author Contribution

J.F.K and B.D. conceived and designed the study. J.F.K, G.L., J.R., N.G., L.L., and C.S. performed the experiments. J.F.K analyzed the data and performed the statistical analyses, with help from V.G. and K.F. J.F.K and B.D. wrote the manuscript with input from J.V., V.G. and K.F.

## Competing Interests

The authors declare no competing interest.

## Acknowledgements

We thank Drs Luca Pinello, Max Trauernicht, Marco Osterwalder, Lucas Ferreira DaSilva, Simon Senan, Michael Love, and Jon Rosen, for valuable feedback on the manuscript and data analysis. We thank the EPFL Flow Cytometry and Gene Expression core facilities, as well as EPFL’s Scientific IT and Application Support (SCITAS). This work was supported by a Swiss National Science Foundation grant (no. 310030_197082), EPFL’s Center for Imaging, a Marie Skłodowska-Curie fellowship (no. 895426), as well as an EMBO long-term fellowship for J.F.K. (1139-2019).

## Methods

### RMCE Cell line establishment

A 3 kb region spanning part of the AXIN2 5’ UTR, the promoter and up to the rs143348853 indel-containing enhancer was targeted by CRISPR-based genome editing using two guide RNAs, of which one is located within the 5’UTR and the other one upstream of the enhancer. The donor construct was synthesized in house and consisted of two homology arms of around 800 bp each, and a loxP – CMV - mCherry - T2A - puromycin resistance - WPRE - bGHpoly(A) – lox2272 - insert cassette. gRNAs and donor cassette and the CAS9 protein were placed on the same plasmid as described previously^2^. Five million MEC1 cells (representing a homozygous REF background) were nucleofected with the CRISPR-donor plasmid using the NEON electroporation system 100 ul kit (Thermo Fisher) with the following settings (3 pulses, 1200V, for 20ms). Cells were cultured with IMDM GlutaMAX with 15% FBS without antibiotics for 48h, followed by selection with 0.75 μg/ml puromycin. After 1-2 weeks, mCherry+ cells were sorted by FACS (Flow Cytometry Core Facility, EPFL) using FACSAriaII or FACSAriaFusion flow cytometers (BD Biosciences) and dispensed as single cells into 384-well plates containing conditioned medium (50:50 fresh 10% FBS, 1% P/S IMDM media with 20% FBS, 0.45 μm filtered medium of cultured MEC1 cells) supplemented with 0.75 μg/ml puromycin; plates were sealed with parafilm and incubated for another two weeks.

Single cell clones were PCR genotyped using three distinct primer combinations. First, only clones that produced a band when using a primer pair targeting the left homology arm and an insert-specific sequence containing the loxP site and an EcoRI motif introduced during cloning. Of the 48 clones selected in the first step, 46 also produced a band when using primers targeting the CMV-mCherry cassette and the right homology arm, confirming integration of both ends. To identify heterozygous clones (one wildtype allele and one CRISRP-modified one), primers targeting both homology arms were used. Only clones that contained two bands (a shorter one representing the wild type allele and a larger one representing the additionally integrated mCherry-puromycin cassette allele) were selected. Lastly, the remaining clones were screened for CRISPR plasmid backbone contamination to guarantee the absence of whole plasmid integration events. For two final clones, the homology-loxP/lox2272 junctions were confirmed with Sanger Sequencing. Both sequencing-confirmed clones were infected with lentivirus expressing Cre recombinase and a blasticidin resistance cassette. The pLV-Cre plasmid was a generous gift from Dr. Jiahuai Han from Xiamen University. The resulting cells were selected with 5 μg/ml blasticidin for 2 weeks and frozen in batches. Due to an insufficient signal strength for sorting mCherry-negative cells from mono-allelic mCherry cells, the mCherry cassette was replaced with a loxP – CMV-GFP-WPRE - bGHpoly(A) – lox2272 (CMV-GFP for simplicity) cassette using CRE-recombinase mediated cassette exchange. The lox-flanked CMV-GFP construct was created by PCR using overhang primers adding lox sites and a lab stock of a CMV-GFP plasmid. The PCR product was inserted into the TOPO vector using the TOPO-blunt-end cloning kit (Thermo Fisher). The resulting vector was nucleofected into one of the plV-CRE / loxP-CMV-mCherry-lox2272 lines and after 1-2 weeks GFP-positive and mCherry-negative cells were sorted by FACS as described in the section before. Single cell clones containing a successfully integrated CMV-GFP cassette were verified by PCR and Sanger sequencing using a similar screening approach as described before, verifying the complete integration of the CMV-GFP cassette with outer/inside and outer/outer primer combinations. A verified clone was grown up, subjected to a bulk FACS sort to select for cells with high GFP signal (top 50% GFP signal), grown for another 3-4 days and frozen in batches of 3-5 Mio cells to serve as a starting point for each replacement experiment.

### Library Synthesis

Construction of long artificial libraries was done using the following approach: individual manipulations within the AXIN2 enhancers or the promoter were introduced by purchasing primers that contained the manipulation as well as an overlap sequence with the sequence right up- or downstream of the manipulation site. The AXIN2 region within the replacement cassette was then amplified as two separate PCR products (up- and downstream of the manipulation site). PCR products were purified by PAGE followed by a clean and concentrator column (Zymo Research). To join two separate PCR products to one long piece, 25-27 cycles of a template-free PCR (TPCR) were performed using a final concentration of ∼15-30nM for each PCR product in a 25ul reaction. The Q5 polymerase part of the NEBNext 2xMasterMix (NEB) was used with the following cycling settings (initial cycle: 3min at 95C, 30s at 55C, 1min at 72C; four cycles using 30s at 98C, 1min at 69C, 1min at 72C, ten cycles using 30s at 98C, 30s at 65C, 1min45s at 72C; ten cycles using 30s at 98C, 30s at 63C, 30s at 72C; final extension for 5min at 72C). Each TPCR was subjected to a final ‘Finish PCR’ (FPCR; Takara ExPremier Polymerase MasterMix; annealing at 60C and extension at 68C), with primers adding i) target-vector overhang for Gibson assembly, a loxP site, a random barcode and a fixed primer sites on the 5’UTR end, and ii) a target-vector overhang, and lox2272 site on the other end. FPCR products were PAGE purified and subjected to another 8cycles of PCR using ‘outer’ (OPCR) primers targeting the target-vector overlap to achieve optimal signal-to-background levels for cloning (Takara ExPremier Polymerase MasterMix). OPCR products were PAGE and column-purified to obtain contamination-free ∼ 3kb long multiplexed inserts.

### Library Cloning

PCR product library pools were inserted into the target vector using Gibson Assembly. For convenience, we used the STARR-seq vector backbone already available in the lab as the target vector as it included a ccdB excision cassette for negative selection. For the initial BC to enhancer mapping experiments (see **Figure 1**), we used the TOPO-XL cloning kit (Invitrogen), however we did not achieve sufficient cloning efficiencies for more diverse libraries. Gibson Assembly products were diluted 1:4 and 2ul of the diluted product was transfected without purification into 50 ul of chemically competent OneShot OmniMax T21R (Invitrogen) chemically competent cells. Cells were incubated on ice for 25 min, heat shocked at 42C for 45s, followed by another 2min on ice, before rescue in 250ul SOC (provided with cells). Cells were incubated at 37C for 1h using a Thermomixer set to 1000rpm before plating on agarose Ampicillin plates. Plates were incubated at 32C for 16-18h followed by another 6h at RT to allow for equally sized colonies. Depending on cloning efficiency and library size, multiple transformations were set up in parallel. Typically, one 2ul transformation of a 3kb fragment resulted in 800-5000 colonies per plate. We noticed that efficiency is sequence and size dependent, with the G/C-rich stretch removal resulting in more colonies compared to the wildtype sequence. It is thus recommended to keep PCR products that differ substantially from each other separate during cloning to allow for a balanced final pool. To provide a specific example, when cloning the library containing modifications to both the *AXIN2* promoter and the enhancer, enhancer overhang primers were mixed prior to PCR amplification, however each distinct promoter manipulation was kept separate. This was done to prevent the G/C-stretch deletion fragments to be overrepresented in the final library pool. To obtain a balanced library, roughly equal numbers of colonies containing a specific construct were mixed (number of plates per unique modification = final library size divided by the number of fragments and the number of colonies per plate) by adding a small volume (∼ 5ml) of LB (with 50 ug/ml Ampicillin) to each plate, scraping off the colonies and combining everything in a round-bottom flask at a final liquid volume of 200 ml. Cultures were incubated at 37C and 190 rpm for a maximum of 2h before extracting the plasmid pool using a Maxi Prep kit (Invitrogen).

### Library transfection and selection of positive cells

The cell line containing the Cre-Blasticidin and the CMV-GFP cassettes were cultured for 3-5 days in IMDM media (Gibco, with 10% FBS and 1% Penicilin/Streptomycin) followed by 5-8 days with the addition of Blasticidin (5ug/ml) to enrich for Cre-expressing cells. At the day of nucleofection, cells were spun down for 5min at 500 g and washed once with Phosphate-buffered Saline (PBS; Gibco). Depending on library size 2 to 6 independent nucleofections were set up. Per reaction, 25-30ug of plasmid library was added to around 5-6 Mio cells before resuspension in 100 ul R buffer. For control nucleofections, no plasmid was added. Cells were nucleofected using the 100ul kit of a Neon Transfection System (Thermo Fisher) with the following settings: 3 pulses of 20ms at 1200V. Cells were immediately transferred to 10ml of antibiotic free IMDM media supplemented with 15% FBS and incubated for 1-2 days before continuing in regular culture medium. Cells were grown up to around 100-150 Mio total cells (7-12 days) for FACS sorting. If FACS was more than 7 days after nucleofection, parts of the cell suspension was discarded. It is critical to thoroughly mix the cell suspension in this case to guarantee a diverse library. On the day of FACS, cells were washed with PBS, resuspended in Fluorobrite with 3% FBS at a final concentration of ∼10Mio cells/ml and filtered. Control cells (no plasmid library) were processed the same way. Library-containing cells were selected by sorting for GFP-negative cells. Depending on library complexity, a total of 200k – 2Mio cells were sorted, and subsequently expanded to around 40-50 Mio cells. From this stock 3-5 replicates of 5 Mio cells each were taken and the remainder of the cell mix was cryo-preserved for later analyses. It is critical to mix the cell suspension sufficiently to optimize equal coverage of barcodes across replicates. This is especially true for suspension cells that form large clumps.

### Estimating Replacement efficiency

For each RMCE experiment a control experiment was included, in which all steps during RMCE were followed, however, using an empty control (no plasmid) during the library nucleofection step. At the beginning of each sort, 100 k control cells and 100k library-replaced cells were sorted and the fractions of GFP-negative cells were compared between the two groups. The fraction of GFP-negative cells in control cells varied between 0.5-2.5% and in library-replaced cells between 1.5-8.9%, with larger fractions in control cells tracking with larger fractions in experimental cells. Typically, a 1.5-6% increase in GFP-negative cells was observed.

### Library extraction, preparation, and sequencing

For each replicate genomic DNA (gDNA) and mRNA were extracted from a total of five million cells using the mini AllPrep DNA/RNA extraction kit (Qiagen). mRNA was purified using the Dynabeads polyA mRNA purification kit (Thermo Fisher). Note, the mRNA purification step is not necessary, however, doing so reduces the number of cDNA reactions required and results in slightly higher mRNA capture rates. The total amount of purified mRNA (typically 0.4-0.8 ug) or total RNA (∼ 30 ug) was reverse transcribed using a mix of three primers targeting the beginning of the second exon of the *AXIN2* gene. For each *μ*l of Maxima H minus RT (Thermo Fisher) up to 0.5*μ*g of poly-A mRNA or up to 3 ug of total RNA was used as input. The resulting cDNA was purified with a Clean&Concentrator kit (Zymo) using 7x binding buffer prior to final amplification. To amplify spliced mRNA, a fourth (closer to the splice-junction) primer targeting the second exon of the *AXIN2* gene, and a library-specific reverse primer placed within the 5’UTR during library synthesis (**Supplemental Figure 1a**) was used and PCRs were run for 16-19 cycles. To determine optimal cycle numbers and correct sizing of fragments a test PCR was run at 16,20,24,28 cycles and evaluated using gel electrophoresis. Final library preparation cycle numbers were chosen to be ∼5 cycles below the cycle producing a faint visible band on the gel (qPCRs failed, likely given the very high GC content). After initial amplification, Illumina adapter sequences and barcodes were added in two separate PCR reactions using 8 and 7 cycles respectively. DNA was cleaned up between each amplification step using magnetic beads (Ampure XP beads). For the final clean-up a double-sided size selection was performed to enrich for the specific lengths of mRNA and gDNA fragments respectively.

Libraries were sequenced at EPFL’s Gene expression Core using a MiSeq or NextSeq desktop sequencer as either single- or paired-end runs depending on availability.

### Barcode to fragment mapping

Library plasmids were digested with restriction enzymes (e.g. AgeI and SalI or BamHI when integrated within the STARR-seq vector) and the resulting DNA was gel purified before ligating library-specific Oxford Nanopore Technologies (ONT) barcodes (SQK-NBD114.24) and adaptor sequences for ONT sequencing. Libraries were sequenced on a Minion or a PromethION 2 Solo device using MINFlow114 and FLO-PRO114M flowcells respectively. The raw fast5 and/or pod5 data files were processed using the guppy basecaller^57^ v6.5.7 provided by ONT on a GPU server. Base called reads were aligned using minimap2 v. 2.24-r1122 on custom reference fasta files containing the different sequences included in each library. Multimappers were removed using samtools v.1.9 such that each read was mapped uniquely to one library sequence. To guarantee accurate mapping, for each read, barcode sequences and uniquely modified region(s) were extracted from the aligned bam files. Further filtering retained only barcodes where the extracted sequences matched unique parts of the actual modification (e.g. for promoter modification) or contained sub sequences mapping to the correct TFBSs (e.g. for synthetic enhancers). For instance, for the 6-GC-box motif deletion, we required sequences to contain exact matches to both the first GC-box and the third box mutation, whereas the two-GC-box deletion had to contain wild type GC-boxes in the third motif location. Only barcodes were retained that were either associated with a single construct or with a mapping frequency at least three-times more than the second best.

### Count data analysis

BCs for both short-read mRNA and gDNA libraries were counted and unique constructs assigned using the long read sequencing data. To plot the raw barcode-level activities, count tables were filtered by requiring that BCs had to be detected in all replicates for the gDNA libraries. gDNA and mRNA counts for each barcode were summed across replicates respectively before computing a BC-level raw activity defined as the mRNA over gDNA ratio. To compute the fraction of expressed barcodes per replicate, we counted the number of barcodes with non-zero counts in the mRNA library and divided it by the total number of BCs detected in the gDNA for a given construct.

To compute average EXTRA-seq activities, we filtered count tables the following way: only BCs with at least 10 total counts across all gDNA replicates were retained and non-zero mRNA counts associated with a zero gDNA count were set to 0. Next, we aggregated all barcodes belonging to a given construct for each replicate separately and computed the log2 fold change using the mpralm function of the mpra package^50^. We chose this package as it estimates the mean-variance relationship (i.e. it corrects for general trends observed between mRNA over gDNA ratio and gDNA counts). To display inter-replicate variability (error bars for each logFC value), we extracted the scaled standard error from the ‘treat’ function of the ‘limma’ package associated with each fit^51^. Normalization to facilitate comparisons across different tools are simply done by subtracting each logFC value by the genotype chosen for normalization (e.g. the REF genotype).

### Puffin model predictions

To obtain predictions based on the machine-learning model Puffin^33^, we used 750bp of sequence up and downstream of the *AXIN2* TSS (excluding the BC) for the wild type and each of the eight promoter modification constructs. Predictions were made for FANTOM-CAGE using the webserver^62^. To compute total expression from Puffin predictions of CAGE, we summed across the 100bp up- and downstream of the INR motif, transformed to log scale and represented everything relative to wild type levels.

### Enformer predictions

To predict expression based on Enformer, we extended the ∼2-3kb of sequence contained within the various library constructs (containing both lox sites, the BC and the library primer site) with 200 kilobases of up- and downstream endogenous sequence directly flanking the landing pad in the *AXIN2* locus. We then trimmed the sequences to a final length of 393,216 bp to conform with the input format for Enformer. Trimming was done in a way that assures that the UTR and promoter sequence is contained within the same bin for all constructs (alignment based on the TSS-downstream side of the sequence). To minimize biases driven by the sensitivity of Enformer to the absolute positional encoding within the input sequence, we used 10 randomly sampled shifted input sequence frames, with shifts being contained within one output bin length (128bp). To mimic the actual experiment, we generated 3-5 random barcodes (7 when systematically assessing the impact different positional encodings have on REF/ALT effect size predictions), to replace the randomized barcode sequence present in our actual libraries. To avoid UTR-sequence specific confounders, the same set of barcodes was used for each construct within a library. This resulted in 30-50 unique predictions for each fragment. Enformer was run using the human set and heads 69, 688, and 5110 representing DNase, CAGE, and H3K27ac levels for GM12878. To compute the total level of expression driven by each construct at the UTR, we first took the log2 of the summed predictions for the three output bins containing, and right up- and downstream of the TSS, before averaging across barcodes and input positional encodings. We did the same to obtain predictions locally at the enhancer, however we used three up and downstream bins of the respective enhancer sequence to capture the entirety of enhancer activity. Importantly, we did not consider bins downstream of the UTR (i.e. there are two alternative TSSs for *AXIN2*, but EXTRA-seq only measures expression stemming from the first TSS and correctly spliced transcripts, see also **Supplemental Figure 1a**). The standard-deviation across individual log-scale predictions (across BCs and positional encodings for a given genotype) are used to represent error bars.

### Local enhancer activity measures

For the comparison of EXTRA-seq enhancer activity with different chromatin marks, accessibility levels, and coactivator levels at enhancers in their native context, fastq files for ATAC-seq, or H3K4me1, H3K27ac, p300, and BRD4 ChIP-seq experiments in GM12878 cells were retrieved from ENCODE (Experiment: ENCSR637XSC for ATAC-seq, ENCFF000ASM, and ENCFF000ATK for H3K4me1, ENCFF000ASP, ENCFF000ASU for H3K27ac, ENCFF000WAM, ENCFF000WAO, ENCFF000WAV, ENCFF000WAX for EP300) or the SRA archive (SRR1636861 for BRD4) and processed as follows: Where applicable, replicates, and read1 and read2 fastq files were combined before aligning to the hg19 reference genome (GRCh37, release 75 from Ensembl) using the bwa-mem tool^63^ with standard settings. Alignment files were sorted and indexed with samtools (version 1.9)^64^ and subsequently transformed to the ‘bigwig’ format using the bamCoverage (version 3.5.0) command from deepTools^43^. To obtain an accessibility value for each enhancer, counts falling within the 200bp used for each endogenous enhancer fragment were summed. For ChIP-seq experiments the 200bp window was extended by 300bp to each side to capture up to two nucleosomes. Summed counts were log-transformed (log2; adding a pseudo count of 1) and normalized by the respective fragment with lowest activity levels.

### STARR-seq

For the STARR-seq experiment of the modified *AXIN2* enhancer, we amplified ∼650bp of enhancer sequence (containing the rs143348853 variant) from the EXTRA-seq library containing the 21 BS variations using PCR primers that add a 12bp unique barcode (towards the 3-prime end), Nextera sequencing adapters, as well as overhangs for assembly into the STARR-seq vector (Addgene #99296). Fragments were cloned into the STARR-seq vector by Gibson Assembly (NEB) and transformed into Dh5A high-efficiency competent cells (NEB). Libraries with an estimated 200 BCs per fragment were generated as described for the EXTRA-seq procedure. Transfections into MEC1 cells were performed in duplicate using ∼6*10^6^ cells and 30 *μ*g STARR-seq plasmid per replicate. Transfections were carried out with the Neon Transfection System (Thermo Fisher) as follows: 3 pulses of 20 ms at 1200 V in R buffer using 100 ul tips. Cells were harvested 24h post transfection and lysed in Trizol (Thermo Fisher). RNA was extracted by chloroform extraction, followed by isopropanol precipitation with a 1:1 ratio. RNA pellets were resuspended in RNAse-free water and digested with DNAseI for 20 min at room-temperature (Zymo). DNAseI-treated RNA was purified with an RNA-clean&concentrator-25 kit (Zymo). A total of 10-12 *μ*g RNA was reverse transcribed using the STARR-seq specific RT primer. For each *μ*l of Maxima H minus RT (Thermo Fisher), 3 *μ*g input was used. RT reactions were diluted 1:4 with RNAse & DNAse-free water before further amplification. The splice-junction PCR was run using the STARR-seq thio-splice-junction primer and a custom reverse primer spanning the SalI restriction site used for linearizing the STARR-seq vector and the read1 Nextera adapter sequence inserted with the library. About one third of the RT reaction was amplified for 20 cycles. Illumina barcodes were added by a final PCR with 8 cycles using the Illumina Nextera guidelines. After each amplification step, PCR products were cleaned up using AMPure XP magnetic beads (Beckman Coulter). Plasmid input libraries were generated by amplifying the plasmid library directly with Nextera Read1 and Read2 indexing primers for 8 cycles (using a total of 600 ng plasmid in 6 separate PCR reactions). Libraries were sequenced either on the same MiSeq run or separately on a NextSeq desktop sequencer at the same facility. To map barcodes to the individual BS variations, the STARR-seq vector was linearized and the fragments sequenced without amplification on the ONT Minion device. Each read was then scanned for an exact match to one of the 21 BS variations and linked to its downstream barcode. Barcodes mapping to more than one BS were discarded from further analyses. Average STARR-seq activities across all barcodes associated with a given enhancer genotype were obtained by following the same procedure as described for the EXTRA-seq setup, with one difference: Instead of gDNA counts, the plasmid DNA counts were used to compute a final log fold change in RNA over DNA expression using the mpralm package.

### Alphafold predictions

We used the alphafold^3^ webserver (https://alphafoldserver.com) using the forward and reverse strand of either the ‘BC11A’ or ‘PU1’ top TFBS modified *AXIN2* enhancer DNA seqeunce, extending a few bps up and downstream of the inserted monomeric and the existing dimeric ETS sites respectively. In addition, we included three full length protein sequences of the TF PU1 (UniProt ID: P17947). Predictions were run using default settings and the top model was used for visualizations using PyMOL.

### Comparison between wildtype and TATA-box promoters

For the comparison between synthetic enhancer activities in combination with either the wildtype or TATA-box insertion promoters as a function of the number of TFBSs, we normalized all genotype-specific logFC values by the REF logFC first. Synthetic enhancers containing a CTCF site were removed from the analysis given CTCF’s ability to serve as a boundary element, thus running counter the idea of classical enhancer function.

### Integrating STARR-seq and Enformer predictions

To combine STARR-seq and Enformer-derived predictions of enhancer activity, we summed the log-transformed and normalized enhancer activities. Normalization was done using the enhancer deletion construct (wildtype promoter) as it was included in all tools. For each combination, the log-scale STARR-seq activities were added with one of the four Enformer-based predictions (log-scale CAGE at the TSS, or CAGE, DNAse, and H3K27ac at the enhancer, see aforementioned section).

### Statistics & reproducibility

All statistical analyses were carried out in R and figures were generated using the ggplot2 package (version 3.4.2). Raw p-values are visualized as ns = p > 0.05, * = p < 0.05, ** p < 0.01, *** = p < 0.001, **** = p < 0.0001. P-values < 2.2*10^−16^ are reported as ‘<< 0.0001’. P-values are unadjusted unless otherwise indicated. Statistical significance between groups was assessed using an unpaired two-sided Mann-Whitney test unless otherwise specified.

## Code and Data availability

Raw and processed sequencing data for STARR-seq and EXTRA-seq experiments, as well as processed summary statistics will be made available upon publication. Custom code will be deposited to GitHub upon publication and is available upon request.

## Notes

### Competing Interest Statement

The authors have declared no competing interest.

