## Supplemental Figures for "EXTRA-seq: a genome-integrated extended massively parallel reporter assay to quantify enhancer-promoter communication"

**a** EXTRA-seq *AXIN2* 5'UTR modification:

```
TCACTGGCTCCGCGAGCCTGGCCCGGGGGAGTCGGCTGGAGCCGGCTGCGCTTTGATAAGGTCCTGGCAACTCAGTAACAGCCCGAGAG
CCGGGAAATAAAAATAACCCCTCAGCATGACGGACAGACC[BC]ATAACTTCGTATAGCATACATTATACGAAGTTATGAGCGATGGAT
TTCGGGGCCGCCCGGCGGCCGAGGCGCCCGCCGAAGGCCCTGCTGTAAAAGGT
```

| | |
|---|---|
| TCACT | INR motif +1 A |
| **CTCAGCATGACGGACAGACC** | fixed primer site for lib amplification |
| **[BC]** | random barcode (10-14 bp) |
| **ATAACTTCGTATAGCATACATTATACGAAGTTAT** | loxP |
| **GT** | 5' splice donor dinucleotide of first *AXIN2* intron |

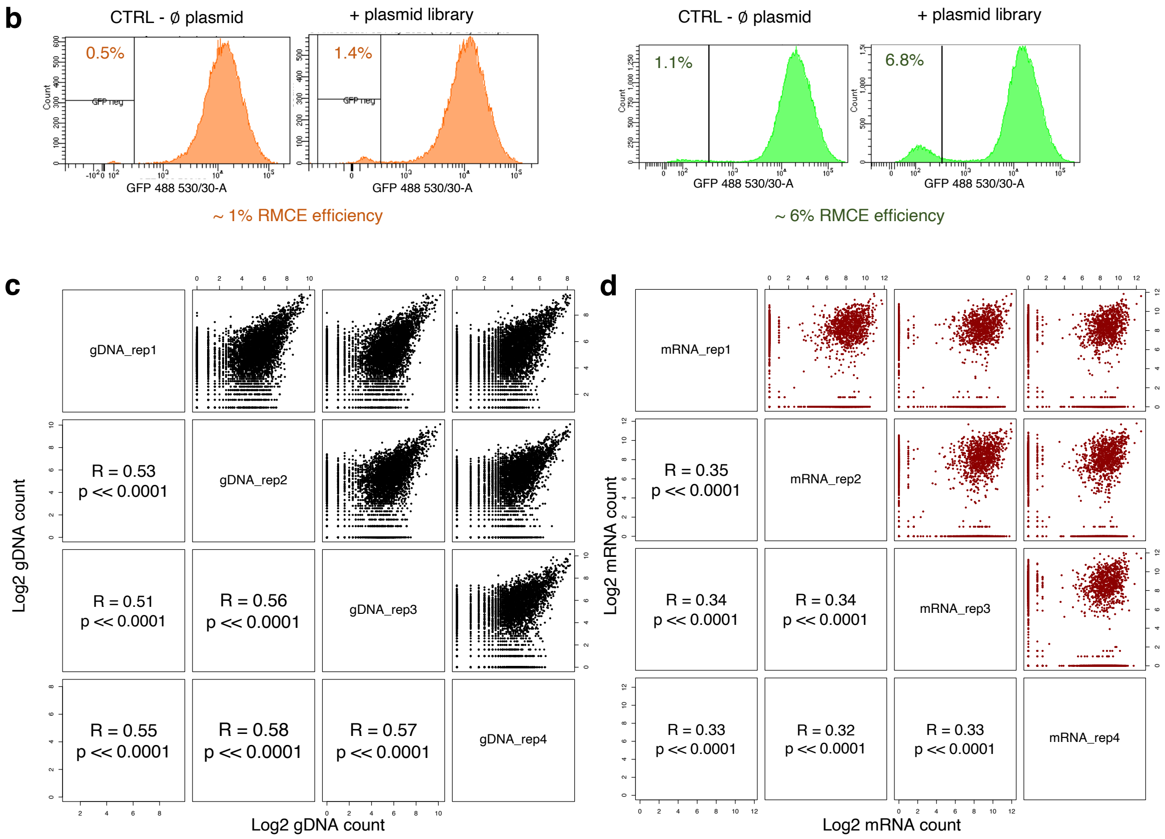


**Supplemental Figure 1 Overview of the *AXIN2* locus and rs143348853 validation experiment**

**a** Shown is the 5'UTR sequence of the *AXIN2* gene including key motifs and the manipulations introduced as part of the EXTRA-seq cassette.
**b** GFP FACS profiles of cells transfected with (right hand side) and without (CTRL = control; left hand side) plasmid libraries. Plots represent lower and upper bound of recombination efficiencies.
**c** Replicate agreement for gDNA counts of individual barcodes for the validation experiment reconstituting either the REF or ALT version of the rs143348853 variant. Correlations and p-values are shown in the lower triangular matrix. P-values < $2*10^{-16}$ are represented as '<<0.0001'.
**d** Same as in c but for mRNA counts.

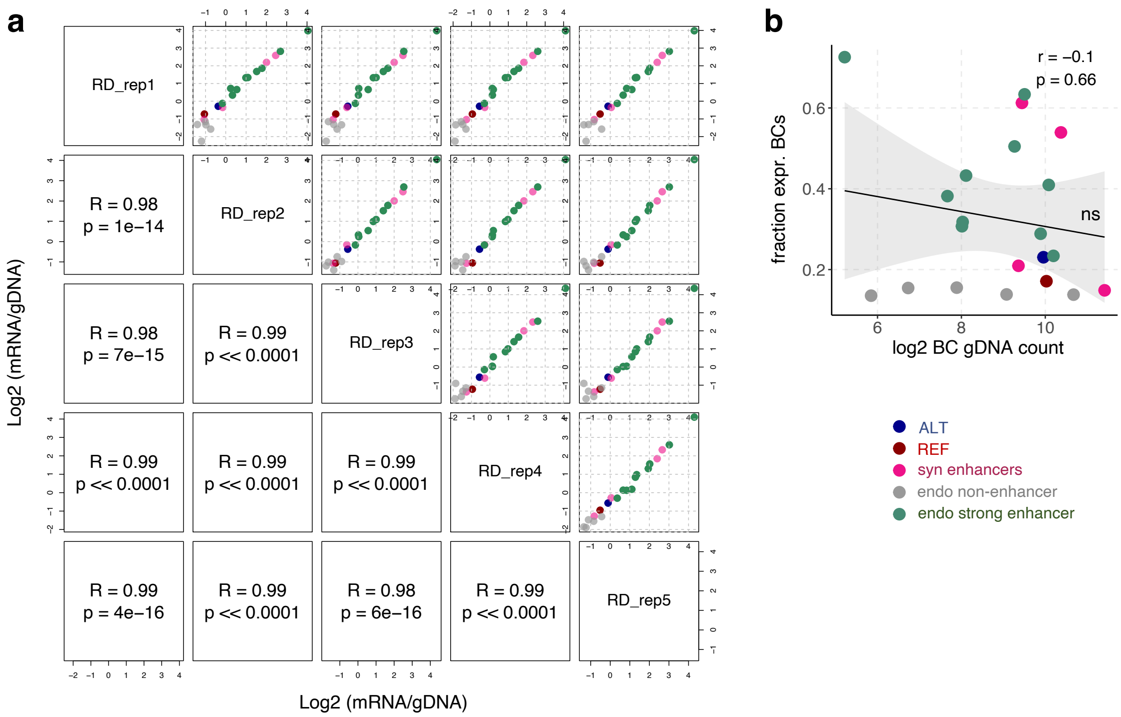


**Supplemental Figure 2 Quality Control for ‘out-of-context’ enhancer library**

**a** EXTRA-seq replicate agreement for aggregate log2 mRNA over gDNA ratios across tested genotypes in the ‘out-of-context’ enhancer library. Pearson correlation values and p-values are shown in the lower triangular matrix. P-values < $2*10^{-16}$ are represented as ‘<<0.0001’.

**b** Absence of relationship between the expression level of a given genotype and its prevalence in the gDNA pool. Fraction of expressed barcodes belonging to a given genotype is shown on the y-axis versus the total log2 count on the x-axis. Pearson correlation value and p-value are indicated in the plot. Straight line represents linear fit.

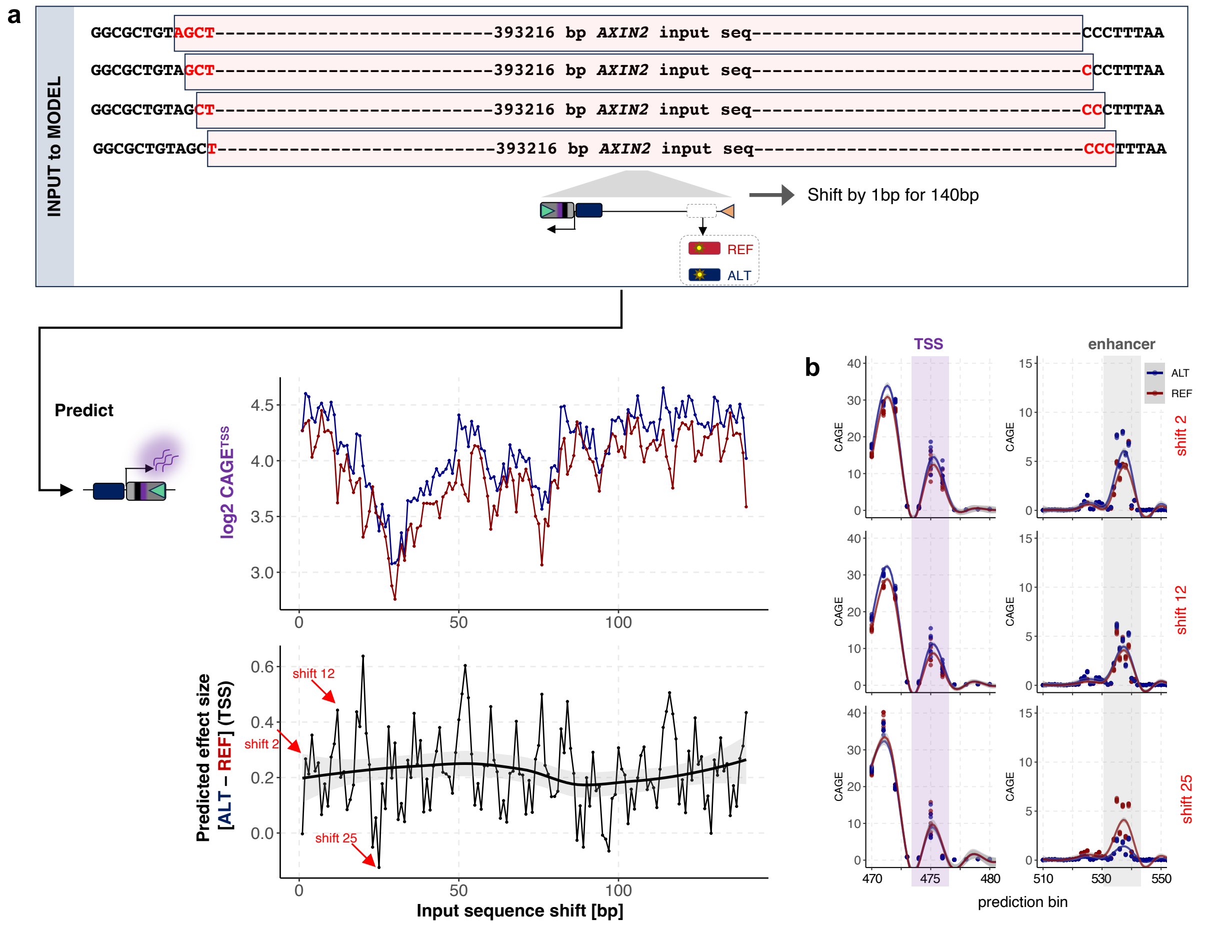


**Supplemental Figure 3 Probing the variability of Enformer predictions on positional encoding within input seqeunce**

**a** Overview of our approach to test Enformer's sensitivity to the exact positional encoding within the almost 400kb of input sequence. The input sequence, containing either the REF (red) or ALT (blue) EXTRA-seq constructs (placed centrally), is shifted 1bp at a time for a total of 140 bps. For each frame, the predicted CAGE signal (log-scale, summarized across three bins) at the TSS is extracted for both genotypes (top line plot). The bottom plot shows the predicted effect size between the REF and ALT genotypes at each shift. Smoothed line represents the average effect size across all shifts (bottom plot). Red arrows and text indicate input sequence shifts for which raw prediction values are shown in b.

**b** Raw Enformer CAGE predictions (y-axis) at the TSS (left) or at the enhancer (right) for both ALT (blue) and REF (red) genotypes for three different input sequence shifts. Bin position is indicated on the x-axis. Points represent predictions for seven random barcodes inserted in the UTR for both genotypes. Lines indicate smoothed values per genotype and across prediction bins. Purple (3 bins) and grey (7 bins) bars indicate the bins used to generate the summarized CAGE predictions shown in **a**.

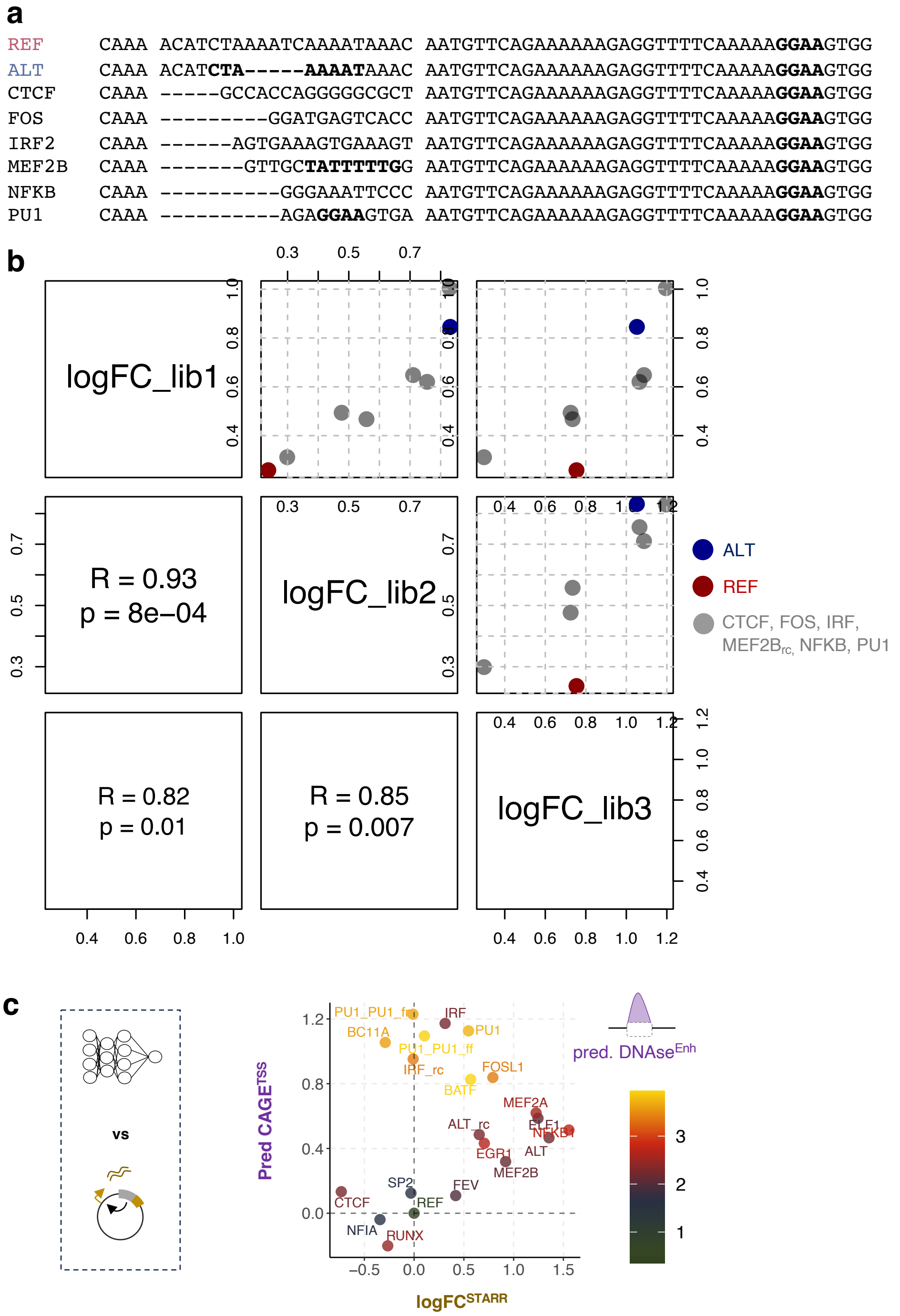


**Supplemental Figure 4 Quality control for single TFBS replacement EXTRA-seq library**

**a** Shown are the TFBSs embedded within the native *AXIN2* enhancer sequence for the eight different enhancer genotypes that were tested in three independent transfections.

**b** EXTRA-seq-measured expression log fold change for the eight enhancer genotypes shown in **a** across three independent library transfections. Lower triangular matrix displays Pearson's correlation values, as well as the associated p-values.

**c** Comparison between Enformer-predicted (y-axis; CAGE at the TSS) and STARR-seq-measured enhancer activity (x-axis) for a total of 21 tested *AXIN2* enhancer genotypes. Point color represents the Enformer-predicted accessibility at the enhancer for each genotype.

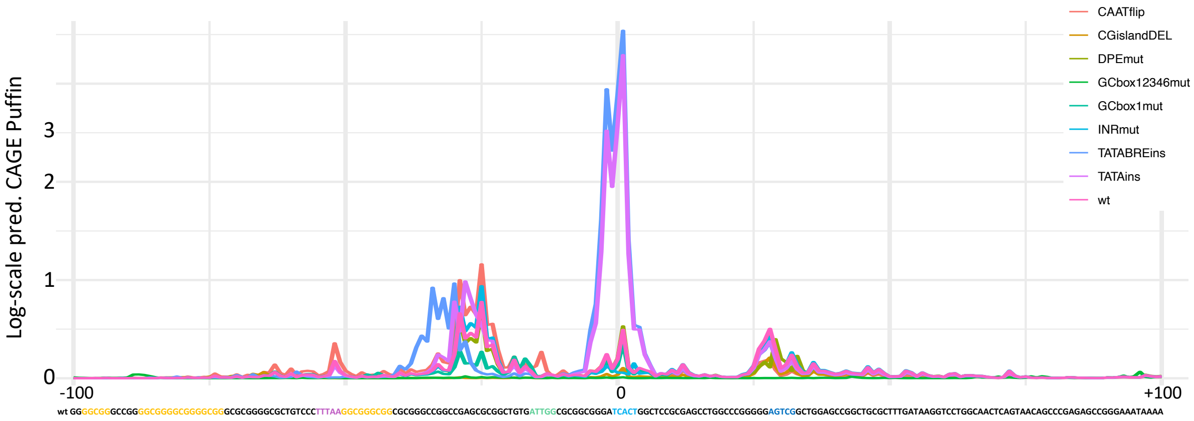


**Supplemental Figure 5 Puffin predictions of FANTOM-CAGE**

Shown are the Puffin model-based predictions in log scale for the nine promoter variations shown in **Figure 4**. The wildtype promoter sequence is shown beneath, with known promoter motifs highlighted in different colors (SP-motif = orange; NFY motif = green; INR motif = cyan; DPE motif = blue). Predictions are centered around the INR motif. The +/– 100bp displayed represents the entire window used to compute the total predicted expression in **Figure 4**.

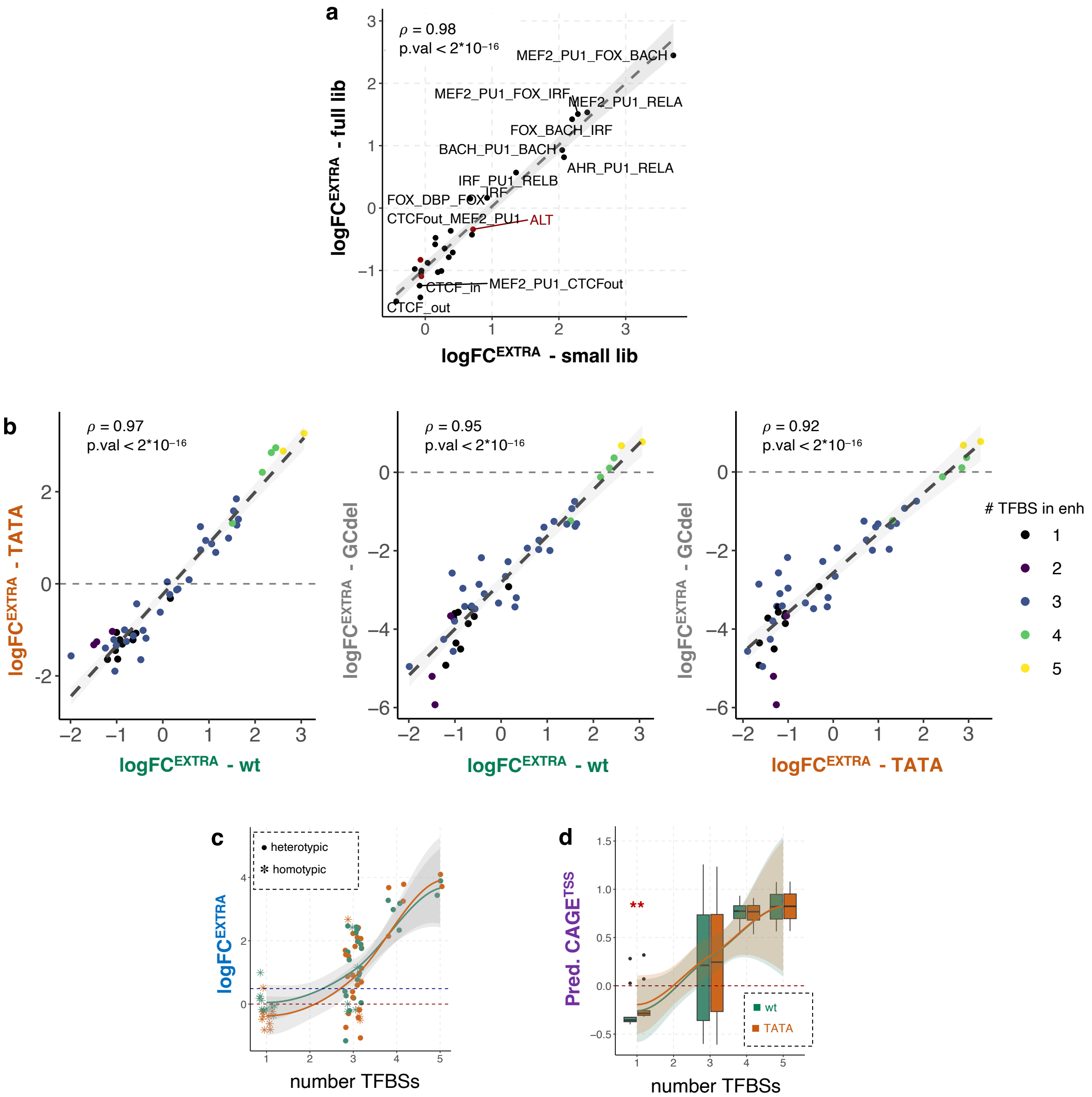


**Supplemental Figure 6 Assessing differences in enhancer activity in function of promoter type**

**a** Comparison of EXTRA-seq-derived enhancer activities (log fold change in expression) for enhancer genotypes included in two separate libraries (small and full library) and independent experimental setups. Pearson's correlation value and associated p-value are indicated above.
**b** Comparison of EXTRA-seq-derived enhancer activities across 47 synthetic enhancer genotypes for the three tested promoter variations (wildtype = wt, TATA-box insertion = TATA, and G/C-stretch-deletion = GCdel). Pearson's correlation value and associated p-values are indicated above, and the total number of TFBSs within each enhancer is indicated by the blue-yellow color palette.
**c** EXTRA-seq-derived enhancer activities split by promoter type (orange = TATA, green = wildtype) and number of TFBSs (x-axis). Enhancers composed of heterotypic or homotypic TFBS combinations are indicated by different shapes.
**d** Enformer-predicted expression (CAGE at the TSS) across synthetic enhancers split by promoter type and the number of included TFBSs. Red stars indicate significant differences in predicted expression levels (TATA > wildtype promoter; Mann-Whitney test; p-value = 0.01).
